# Sulcal landmarks reveal lineage-specific trajectories and multi-scale specialization of the mammalian cortex

**DOI:** 10.1101/2025.10.20.683586

**Authors:** Songyao Zhang, Mingrui Zhuang, Yanan Tian, Weichen Yan, Xi Jiang, Marco Palombo, Tuo Zhang, Hongkai Wang

**Affiliations:** Central Hospital of Dalian University of Technology, Dalian, China; School of Biomedical Engineering, Faculty of Medicine, Dalian University of Technology, Dalian, China; Liaoning Key Laboratory of Integrated Circuit and Biomedical Electronic System; School of Life Science and Technology, MOE Key Lab for Neuroinformation, University of Electronic Science and Technology of China, Chengdu, China; Brain Research Imaging Centre, Cardiff University, Cardiff, Wales, United Kingdom; School of Automation, Northwestern Polytechnical University, Xi’an, China

## Abstract

An important topic of evolutionary neuroscience is to understand how brain function and structure across evolutionary time. In the absence of functional data for ancestral species, studies often rely on structural features such as cortical volume or surface area. However, atlas-based metrics derived from human or a few model organisms lack biological interpretability across species with divergent cortical architectures. Here, we investigate sulcal pits—the locally deepest points of cortical folds—as conserved, atlas-independent landmarks that provide a robust basis for cross-species comparison. Using cortical surface reconstructions from 90 mammalian species, we show that sulcal pit distribution patterns partly recapitulate phylogenetic relationships, independent of overall brain size (volume and surface area), and vary systematically with ecology and lifestyle. By tracing pit-based evolutionary trajectories from 80 million years ago to the present, we found pronounced differences across ecological and behavioral categories, indicating that cortical folding has diversified in close association with species’ habitats, lifestyles, and social structures. Using sulcal pits, we identified *Homo sapiens*-specific regions and species-shared regions. The *Homo*-specific areas were functionally associated with higher cognitive and emotional processes, distinguished by unique histological features, enriched for gene sets related to neural regulation, and exhibited markedly different cell-type abundance profiles compared to shared regions. Together, these results establish sulcal pits as a robust, evolutionarily informative feature for cross-species cortical alignment, offering new insights into the structural innovations underlying brain evolution study.

## Introduction

Comparative analysis across species has long been central to understanding mammalian brain evolution. Advances in non-invasive imaging techniques such as MRI, DTI, and CT have enabled systematic measurement of structural features across species [1]. Global metrics such as brain volume and cortical surface area have been linked to social complexity, tool use, and advanced cognition in primates [2, 3, 4, 5], but these associations provide only a coarse view of evolutionary processes. To move beyond global features, many studies have examined regional evolution using atlas-based frameworks, in which human-defined parcellations are projected onto other species to compute cortical thickness, surface area, or gyrification [6, 7, 8]. While these approaches have revealed species differences in association cortices, they assume consistent regional correspondence across species—a biologically unrealistic premise [9, 10]. Species-specific atlases (e.g., for macaques, mice, and chimpanzees) partly address this issue [11, 12, 13, 14], but they cannot be extended to extinct taxa and depend on large-scale data from extant species.

To overcome these challenges, researchers have increasingly turned to cortical folding morphology, which provides an atlas-independent and biologically grounded descriptor of brain structure. Unlike metrics such as cortical thickness, volume, or surface area, folding reflects the macroscopic expression of cellular processes including proliferation, migration, and differentiation [15]. An increasing body of evidence suggests that folding patterns are not merely byproducts of brain development, but instead play a fundamental role in determining the organization and efficiency of brain function [16]. Distinct folding patterns have been linked to individual differences in cognition, brain network organization, and vulnerability to neuropsychiatric disorders [17, 18]. The convoluted folding of the cerebral cortex represent a key evolutionary adaptation observed across many mammalian species, with species-specific variations highlighting the need for comparative studies of cortical gyrification [19]. From an evolutionary perspective, interspecies differences in folding complexity are thought to reflect variations in cognitive capacity and the number of functional modules, with more extensively folded cortices typically associated with higher-order functional specialization and advanced cognitive abilities [20]. To quantify these differences, researchers have proposed metrics such as the curvature, gyrification index (GI), and fractal dimension (FD) to characterize folding complexity [21, 22, 23]. However, like cortical thickness and surface area, these metrics primarily reflect outcomes of the late-stage folding process, and offer limited insight into the underlying mechanisms that initiate cortical folding [24, 25]. Moreover, these features often exhibit large inter-individual variability within the same species, which reduces their stability and reproducibility in cross-species comparisons and phylogenetic modeling [26].

By contrast, cortical landmarks such as sulcal pits—defined as the deepest points within sulci—are thought to arise at the earliest stages of cortical folding and offer a more developmentally grounded and spatially precise alternative [27, 28]. Sulcal pits have been shown to appear early in development and demonstrate high spatial consistency across individuals of the same species [29]. More importantly, the distribution of pits in the human brain has been shown to parallel the shape of the lateral ventricles, suggesting that pits may serve as a structural template for cortical folding patterns [27]. Unlike atlas-based regional statistics, landmarks provide an atlas-independent framework that can be generalized to any species and require only a small number of subjects per species for reliable estimation [30]. Because sulcal pits are defined purely by cortical morphology, they can be extracted from any dataset that allows reconstruction of cortical surfaces, whether derived from MRI, CT, and potentially even from even fossil endocasts, thereby opening possibilities for extending comparative analyses to extinct taxa. As such, sulcal pits offer a high-resolution, stable, and scalable structural basis for cross-species comparison and for studying brain evolution across species.

In this study, we utilized cortical surface data from 90 mammalian species published in previous studies to construct the largest cross-species sulcal pit distribution database to date. We show that pit patterns recapitulate phylogenetic relationships, and this relationship is independent of overall brain size, including volume and surface area, and that they vary systematically with ecology and lifestyle, revealing constraints related to lineage and environment on cortical folding. By tracing pit accumulation along evolutionary timelines, we uncovered pronounced differences across ecological and behavioral categories and identified species specific evolutionary turning-point times in cortical complexity, with closer relatives tending to show more proximate turning points. Focusing on *Homo sapiens* and its closest relatives, we further delineated *homo*-specific cortical regions enriched for higher-order cognitive and emotional functions, distinguished by unique histological microstructure, enriched for gene programs related to neural regulation, and characterized by divergent cell type abundance profiles. Together, multi level evidence establishes sulcal pits as a feasible and effective medium for cross species cortical comparison and evolutionary inference.

## Results

### Sulcal pit counts are associated with species’ ecology and behavior (Fig. 1)

**Fig. 1A-B** illustrate the process for pit extract, using the *Homo sapiens* and *Ochotona macrotis* as examples. For each species, we first constructed a 3D convex hull of the gray matter surface. As shown in **Fig.1A**, the red closed curve denotes the outer contour of a 2D slice of the cortical surface, whereas the blue closed curve represents the corresponding contour of the convex hull. Sulcal depth at each vertex was defined as the euclidean distance to the nearest point on the convex hull. Local maxima of sulcal depth were then identified to determine sulcal pit locations, which are shown as yellow spheres in **Fig. 1B. Fig. 1C** shows the phylogenetic tree of 90 species analyzed in this study, which included 58 primates, 28 rodents, 2 lagomorphs, 1 scandentia, and 1 dermoptera. To compare differences among ecological and behavioral classifications, we calculated whole-brain pit counts across all 90 species, as illustrated in **Fig. 1D**. Significant differences in pit counts were observed between primates and non-primate (Cohen’s d = 1.78, *P*_FDR_ *<* 0.001), large group and non-large group species (Cohen’s d = 1.15, *P*_FDR_ *<* 0.001), as well as fossorial and non-fossorial species (Cohen’s d = -1.06, *P*_FDR_ *<* 0.001).

**Figure 1.**
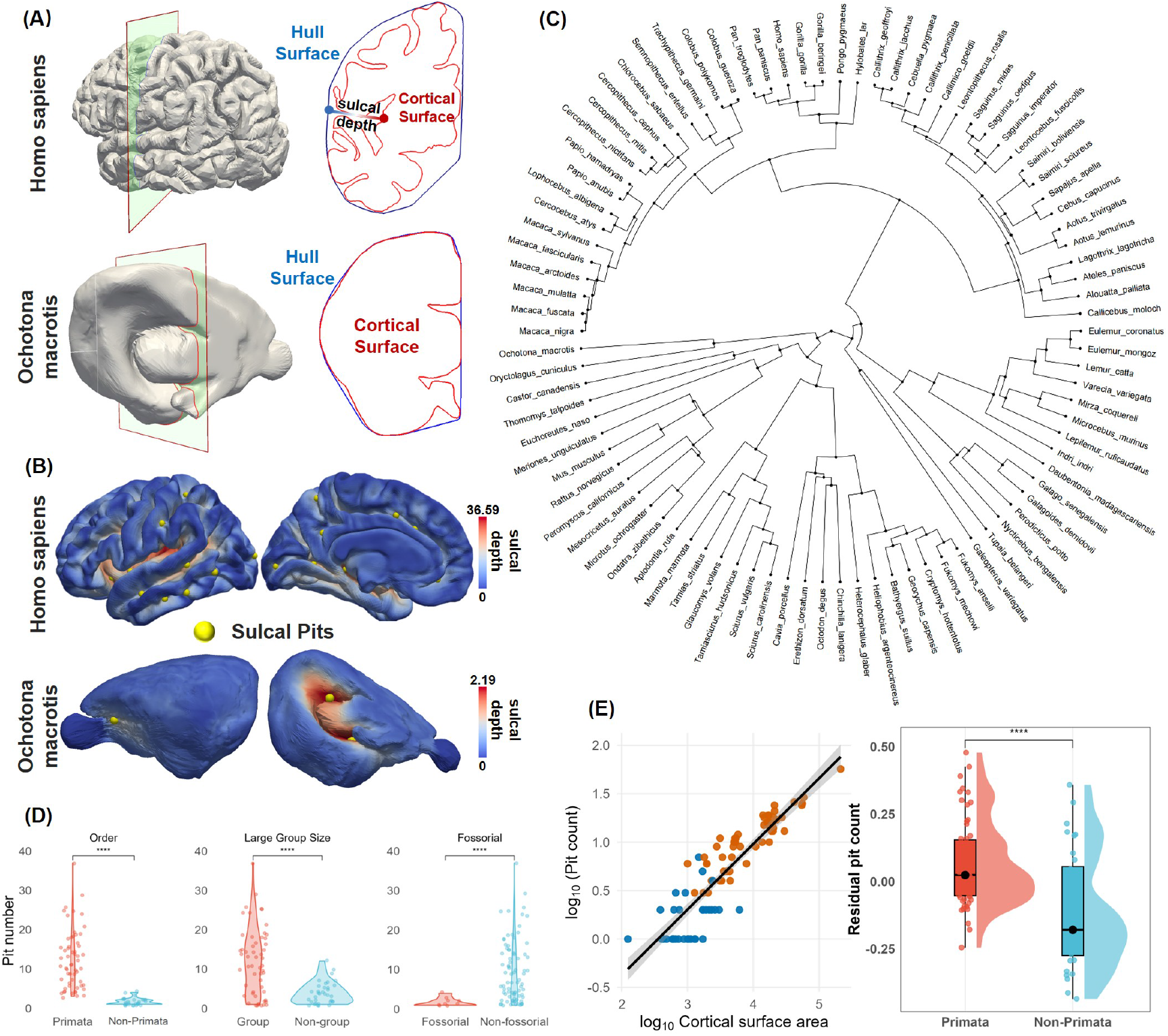
Sulcal pit extraction process and pit count statistics across 90 species. (A) Using the cortex of *Homo sapiens* and *Ochotona macrotis* as examples, the 3D convex hull of the gray matter surface was constructed. The red closed curve represents the boundary of the 2D slice, while the blue contour outlines the corresponding 2D slice of the convex hull. (B) Sulcal pits were identified as local maxima of sulcal depth (yellow spheres). (C) The phylogenetic tree of the dataset used in this study, including 90 species. (D) Whole-brain pit counts stratified by taxonomic order (Primata vs. Rodentia), social organization (large group vs. non-group), and habitat (fossorial vs. non-fossorial). Each point denotes a species, and violins summarize distributions. Significance was assessed using two-sided wilcoxon tests with FDR correction. (E) Left: Log–log scaling of pit count with cortical surface area. Scatter plot of log_10_(pit count) versus log_10_ surface area with an OLS fit and 95% confidence band (points colored by primate vs. non-primate). Right: Residual pit counts after adjusting for cortical surface area, showing a significant difference between primates and non-primates.

Because whole-brain pit counts may covary with brain size, we modeled log_10_(pit count) against log_10_ *A* and log_10_ *V*, where *A* denotes cortical surface area and *V* brain volume. In univariate log–log fits, log_10_(pit count) increased with log_10_ *A* with slope 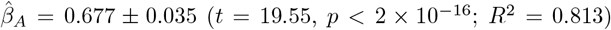, as shown in **Fig. 1**E (left). Interpreted elastically, a tenfold increase in *A* corresponds to ∼ 10^0.677^ ≈ 4.8*×* more pits. The relationship with *V* was similar 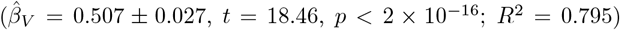, implying ∼ 10^0.507^ ≈ 3.2*×* more pits per tenfold increase in volume (Supplemental Information, Fig.S1). Thus, brain size explains ∼ 79–81% of the log variance in pit number, leaving ∼ 19–21% unaccounted for by area or volume alone. Importantly, in a multivariable log–linear model jointly adjusting for *A* and *V*, size-adjusted (residual) pit counts remained significantly different across categories—including primates vs non-primates (**Fig. 1**E, right), arboreal vs non-arboreal, and terrestrial vs non-terrestrial (Supplemental Information, Fig.S2). Even this global measure—the whole-brain pit count—therefore carries ecology- and behavior-related signal that is not reducible to brain size alone, motivating the more detailed analyses that follow.

### Sulcal pit distribution similarity is associated with phylogenetic similarity among species (Fig. 2)

To examine whether interspecies similarity in sulcal pit distribution partially reflects phylogenetic relatedness from a macroscale anatomical perspective, we computed both a phylogenetic similarity matrix (PS) and a pit distribution similarity matrix (PDS) across species (**Fig. 2A**). The phylogenetic similarity matrix, derived from the evolutionary tree by quantifying pairwise species relationships, exhibited a hierarchical structure: at a coarse level, species clustered into two broad taxonomic groups (pink boxes); at a finer scale, they segregated into five distinct clusters (red boxes, M1–M5), each exhibiting significantly greater within-group than between-group phylogenetic similarity. These five clusters correspond to five suborders: *Haplorrhini, Strepsirrhini, Hystricomorpha, Sciuromorpha*, and *Myomorpha*. The sulcal pit distribution similarity matrix, constructed by quantifying the spatial similarity of pit patterns across all species pairs (details see Methods), revealed a comparable hierarchical structure to that of the phylogenetic similarity matrix. This correspondence suggests that the spatial organization of sulcal pits is, to some extent, shaped by evolutionary lineage. A Mantel test confirmed a significant positive correlation between the two matrices (*r* = 0.35, *p <* 0.001), supporting the notion that species with closer evolutionary relationships tend to exhibit more similar pit distribution patterns (**Fig.2B**, left panel). The correlation was more pronounced within primates (*r* = 0.41, *p <* 0.001) and weaker yet still statistically significant among rodents (*r* = 0.11, *p <* 0.05), indicating lineage-specific variation in the degree of evolutionary constraint on pit organization.

**Figure 2.**
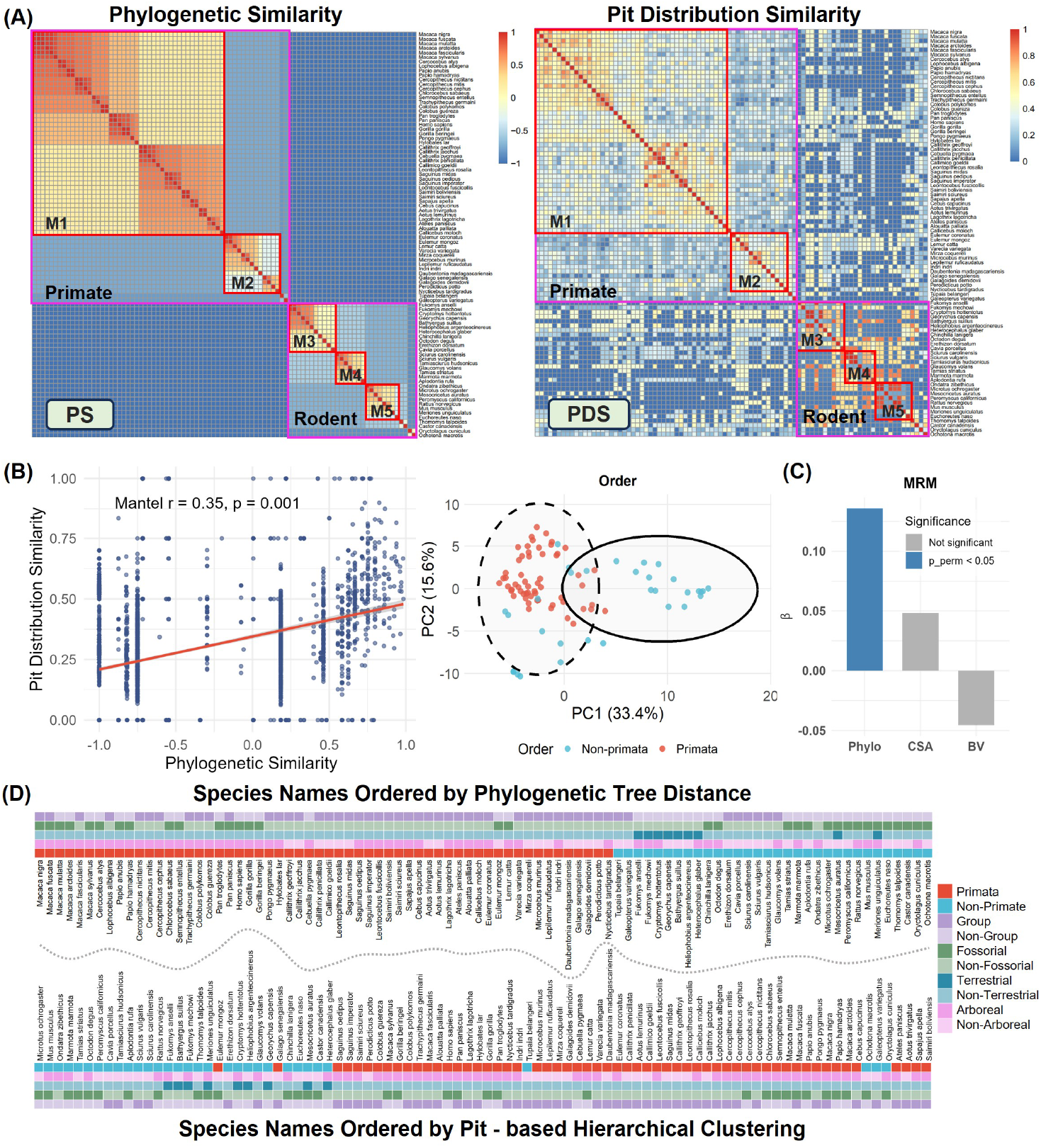
Phylogenetic similarity and pit distribution similarity across species. (A) Left: Phylogenetic similarity matrix derived from the evolutionary tree. Right: Pit distribution similarity matrix, showing a similar structure to the phylogenetic matrix. (B) Left: Relationship between phylogenetic similarity and pit distribution similarity across species pairs. Each point represents a species pair, and the red line indicates the fitted trend. Right: PCA of the phylogenetic similarity matrix. The scatter plot shows species scores on the first two principal components. K-means clustering (*k* = 2) in PC space separates primates (red) and non-primates (blue). (C) Multiple regression on distance matrices (MRM). Standardized coefficients (*β*_std_) for phylogenetic distance, cortical surface area difference, and brain volume difference predicting sulcal-pit dissimilarity. Colors indicate permutation significance (10,000 permutations): blue, *p*_perm_ *<* 0.05; grey, n.s. Only phylogenetic distance was significant (*β*_std_ ≈ 0.136, *p*_perm_ *<* 0.001), whereas area and volume differences were not significant (both *p*_perm_ *>* 0.4). (D) Species ordered by evolutionary tree (top) and hierarchical clustering of pit distribution similarity (bottom).

To assess whether pit-distribution similarity organizes species along a compact set of axes consistent with ecological or phylogenetic structure, we conducted principal component analysis of the similarity matrix. The right panel of **Fig. 2B** shows the first two components explained 33.4% and 15.6% of the variance. K-means clustering (*k* = 2) in this space yielded a clear separation primarily along PC1, aligning largely with primate versus non-primate membership. This indicates that pit-based similarity embeds strong phylogenetic signal in a low-dimensional representation, consistent with ecology- and lineage-related organization. To test whether the correspondence between pit similarity and phylogenetic relatedness could be explained by differences in brain size, we applied multiple regression on distance matrices (MRM) [31], which simultaneously tests the effect of several predictors on pairwise dissimilarity (**Fig.2C**). The model included phylogenetic distance, cortical surface area difference, and brain volume difference as predictors. Consistent with the Mantel results, phylogenetic distance (Phylo) remained the only significant predictor of pit dissimilarity (standardized *β* = 0.136, *p*_perm_ *<* 0.001), whereas cortical surface area (CSA) and brain volume (BV) differences were not significant (both *p*_perm_ *>* 0.4; Supplementary Information, Table S1). These results reinforce that the observed pit–phylogeny correspondence is robust and independent of allometric scaling effects.

To further evaluate whether pit distribution similarity reflects phylogenetic or ecological structure, we compared species orderings derived from the evolutionary tree and from hierarchical clustering of the pit similarity matrix (**Fig. 2D**). Despite the clustering being performed without phylogenetic input, the resulting arrangement broadly separated primates from non-primates, indicating that sulcal pit architecture carries a strong phylogenetic signal. Other ecological or behavioral categories, such as large group size and lifestyle (fossorial, terrestrial, arboreal), did not form equally clear global partitions. This suggests that while pit similarity is primarily constrained by lineage, ecological and lifestyle influences are expressed in more localized or subtle patterns.

### Sulcal pit reveals the laws of brain evolution (Fig. 3)

The process of species evolution involves an ancestral species gradually diverging into two independent species over long periods of geographic isolation or environmental changes, with brain cortical folding complexity also shifting throughout this process. Since pits have been shown to reflect phylogenetic relationships among species, exploring patterns of brain evolution through spatial distribution of sulcal pits provides a valuable research approach.

**Figure 3.**
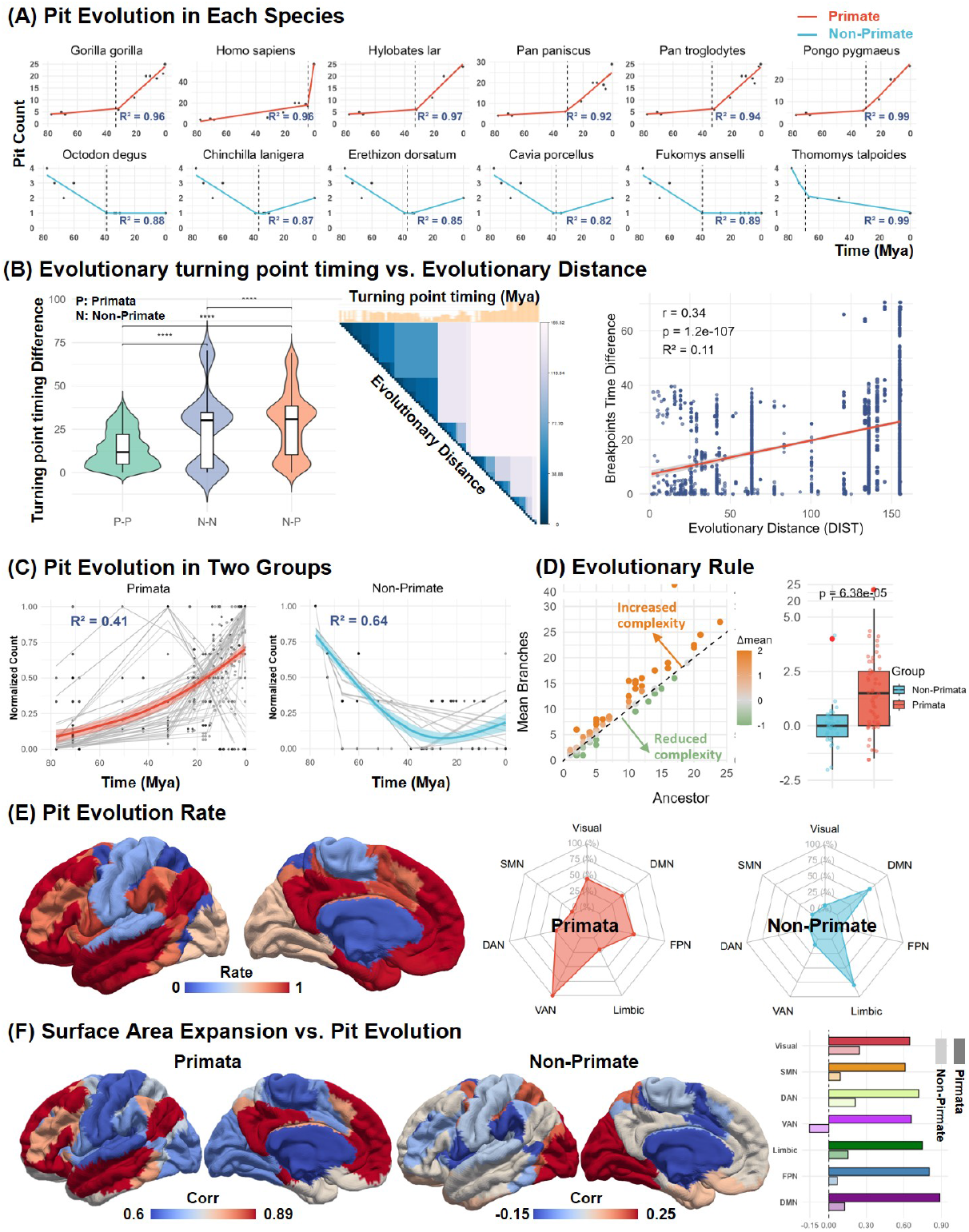
(A) Representative piecewise linear fits of whole-brain pit counts for six randomly selected primate and six non-primate species, with *R*^2^ indicating the goodness of fit. (B) Cross-species similarity in the timing of evolutionary turning points. Left: Violin plot shows the distribution of turning point time differences across species pairs, categorized into three groups based on phylogenetic relationships. These group differences are statistically significant (wilcoxon tests, *p <* 0.001). Center: Pairwise evolutionary distances with corresponding species-level turning point timing above. Right: Scatter plot of evolutionary distance (x-axis) versus turning point timing difference (y-axis) across all species pairs. (C) The evolutionary trajectories of pit counts in primates and non-primates. Gray lines show species-level piecewise fits, and colored curves show group trends estimated with GAMs with 95% confidence intervals. (D) Evolutionary change in pit-defined folding after lineage splits. Left: descendant mean versus ancestor with the *y* = *x* line. Right: boxplots show greater post-split increases in primates than in non-primates (two-sided Wilcoxon, *p* = 6.38 *×* 10^−5^). (E) Left: Growth rate of sulcal pit counts in seven brain networks. Right: Proportion of species in which each network showed the fastest evolutionary rate. (F) Coupling between cortical surface area expansion and sulcal pit growth rate across the seven brain networks.

To quantify temporal change along each lineage, we reconstructed the pit count from the last common ancestor (80 Mya) to each extant species and fit a piecewise linear model to the time series, yielding two phase-specific slopes and a turning-point time (breakpoint). **Fig. 3A** shows representative trajectories from several primates and non-primates to illustrate the metric (early vs late slopes) and turning points (full species panels are provided in the Supplementary Information, Fig.S4). Building on these estimates, we hypothesized that closer phylogenetic relatedness would correspond to smaller differences in turning-point times. We therefore correlated pairwise phylogenetic distance with the absolute difference in turning-point times and summarized the distributions by grouping species pairs as N–N (non-primate vs non-primate), P–P (primate vs primate), and N–P (non-primate vs primate). As shown in the left panel of **Fig. 3B**, the P-P group exhibits the smallest differences in timing of the evolutionary change point, followed by the N-N group, while the N-P group shows the largest differences. The differences among the three groups are statistically significant (pairwise Wilcoxon rank-sum test, *p <* 0.001). This result suggests that species with closer phylogenetic relatedness tend to have more similar timing of the evolutionary change point. To further validate this hypothesis, we analyzed the relationship between phylogenetic distance and differences of turning-point times across all species pairs. The heatmap in the center of **Fig. 3B** shows the evolutionary distances between all species pairs, and the bar plot above displays the turning-point times for each species. The scatter plot on the right displays all species pairs, with the x-axis representing the evolutionary distance between two species and the y-axis indicating the difference in their turning-point times. A positive correlation was observed between these variables (*R*^2^ = 0.11, r = 0.34, p *<* 0.0001), supporting the hypothesis that closer phylogenetic relatedness species tend to have more similar turning-point times.

Aggregating across all species, we then compared group-level evolutionary trends using GAM fits (*k* = 3) for primates and non-primates (**Fig. 3C**). After normalizing within species to [0,1] to remove scale differences, the primate trajectory remains flat for much of deep time and rises steeply over the past 20 Mya, consistent with a late-phase acceleration (GAM *R*^2^ = 0.41). In contrast, non-primates show an early decline (80–30 Mya) followed by a prolonged plateau, yielding a more stable/flattened group trend (GAM *R*^2^ = 0.64). These patterns indicate that the timing and rate of pit accumulation differ across lineages: primates concentrate gains more recently, whereas non-primates change earlier and then stabilize. We repeated the same GAM analysis across ecological groupings (social vs. non-social, diurnal vs. non-diurnal, fossorial vs. non-fossorial, terrestrial vs. non-terrestrial, arboreal vs. non-arboreal; Supplementary Information, Fig.S5). Across these contrasts, pit-based trajectories consistently differentiate groups, indicating that pit-defined evolutionary tempo is sensitive to species’ ecology and lifestyle.

To quantify how cortical folding changes after lineage splits, we compared each ancestor’s pit count with the mean of its two descendants, 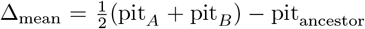 . In the phylogeny from the 80 Mya root to the 90 extant species, there are 89 internal nodes, each providing an ancestor versus descendants comparison. In **Fig. 3D** left, points above the diagonal (*y* = *x*) indicate increased folding complexity in descendants, whereas points below indicate reductions. Most splits fall above the line, with larger positive than negative shifts and decreases typically close to the diagonal. In **Fig. 3D** right, the distribution of Δ_mean_ differs by clade: primates show significantly greater post-split increases than non-primates (two-sided Wilcoxon, *p* = 6.38 *×* 10^−5^). Together, these results indicate that lineage splits tend to yield net gains in pit-defined folding complexity, with the tendency strongest in primates (see also Supplementary Fig.S6 for results using Δ_min_ = min(pit_*A*_, pit_*B*_ ) − pit_ancestor_).

Although sulcal pits were identified in an atlas-independent manner, for cross-species comparison of regional evolutionary speed we summarized pit count changes within the seven canonical functional networks defined in the Yeo 7-network parcellation [32] (**Fig. 3E**). Across 90 species, we tracked trajectories from the last common ancestor to the present. We quantified growth by the increase in pit counts within each functional network. The fastest growing networks were the default mode network (DMN) and ventral attention network (VAN), whereas the slowest were the dorsal attention network (DAN) and somatomotor network (SMN). To further compare the evolutionary pace of brain regions in primates and non-primates, we identified the brain network in each species that showed the largest increase in pit counts from the last common ancestor (80 Mya) to the present. This network was considered the fastest-evolving for that species. We then calculated the proportion of species in which each of the seven networks was identified as the fastest-evolving, separately for primates and non-primates. In primates, the VAN showed the most rapid evolution, followed by the DMN, and the frontoparietal network (FPN). In contrast, non-primates exhibited the fastest evolution in the limbic network, with the DMN ranking second. This contrast highlights distinct selective pressures shaping brain network evolution in primates versus non-primates, potentially reflecting differences in cognitive, perceptual, and emotional processing demands between the two groups.

Previous studies have frequently used cortical surface area expansion as a key indicator of brain evolution, reflecting changes in cortical size across regions and species [33, 1]. While informative, this measure reflects only one dimension of cortical remodeling. Our earlier results show that pits are not simply a proxy for brain size: whole-brain pit counts retain signal after adjusting for area and volume (**Fig. 2E**), and an MRM analysis indicates that the pit–phylogeny association is not driven by size covariation (**Fig. 3**). Therefore, in this section we examine how surface area expansion relates to the evolutionary of folding complexity based on sulcal pits. Specifically, we compute the Pearson correlation coefficient between these two features across functional networks and taxonomic orders (**Fig. 3F**). The results revealed that, in primates, cortical expansion and increased folding complexity exhibited strong synchrony, with PCC values ranging from 0.6 to 0.89. Among the networks, the DMN showed the highest correlation, suggesting that its expansion was accompanied by a substantial increase in folding complexity. In contrast, the somatomotor network (SMN) exhibited the lowest correlation, indicating weaker coupling between surface area growth and complexity increase. In non-primate species, however, the synchrony between cortical expansion and complexity increase was generally weak across all networks, with PCC values ranging from -0.15 to 0.25. The visual network demonstrated the highest synchrony, whereas the VAN showed a negative correlation. Further comparison of the synchrony between surface expansion and folding complexity across the seven networks revealed that the visual network exhibited the smallest difference between primates and non-primates, while the VAN showed the largest disparity. These findings suggest that the co-evolution of cortical expansion and folding complexity is more prominent in primates, potentially reflecting a greater demand for structural complexity in the evolution of advanced cognitive functions.

### Differences in pit distribution between *Homo sapiens* and closely related species reflect their evolutionary strategies across multiple scales (Fig. 4 and Fig. 5)

Building on our previous findings that sulcal pits can reveal evolutionary patterns through differences in cortical folding complexity, we next ask: what cortical morphological changes did *Homo sapiens* undergo to diverge from its closest evolutionary relatives? To address this question, we focus on a comparative analysis of several primate species most closely related to *Homo sapiens*. We selected six species with the closest evolutionary distance to *Homo sapiens* (**Fig. 4A(i)**). These species exhibit cortical morphologies that are more similar to *Homo sapiens* compared to other species. We classified sulcal pits into two categories based on their cortical spatial distribution: *Homo sapiens*-specific and species-shared (**Fig. 4A(ii)**, blue and red spheres; see Methods for details). Although pits were identified in an atlas-independent manner, the classification of Homo sapiens-specific and species-shared regions was derived directly from pit-based analyses. For integrative analyses with brain function, microstructural features, gene expression, and cell abundance, the pit-defined regions were further mapped onto the schaefer-100 parcellation. Within this framework, we identified cortical regions that are predominantly *Homo sapiens*-specific (labeled as -1) or predominantly species-shared (labeled as 1-3, with the value reflecting the degree of sharing).

**Figure 4.**
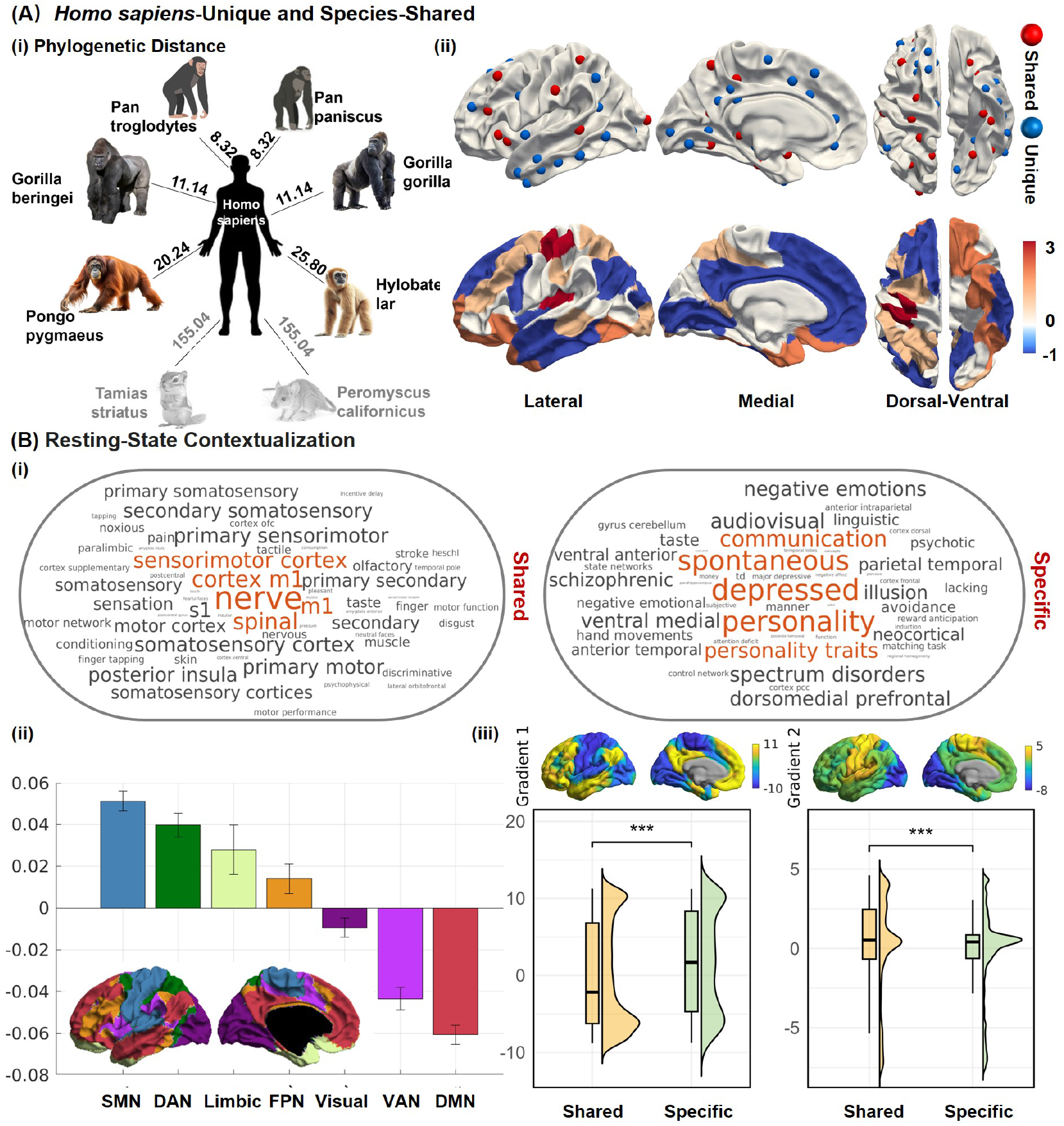
(A) Macroscale functional differences between *Homo sapiens* and closely related species revealed by sulcal pit spatial distribution. (i) Six species with the closest phylogenetic distance to *Homo sapiens* were selected to define *Homo sapiens*-specific and cross-species shared cortical regions. (ii) Red spheres indicate species-shared pits, while blue spheres represent *Homo sapiens*-specific pits. Cortical regions are labeled as predominantly *Homo sapiens*- specific (value = –1) or species-shared (values = 1–3), with higher values indicating a greater degree of cross-species sharing. (B) (i) Meta-analysis of functional annotations associated with species-shared and *Homo sapiens*-specific cortical regions. (ii) Distribution of *Homo sapiens*-specific versus shared sulcal pits across brain networks defined by the Yeo7 functional atlas. (iii) Principal functional gradients (G1 and G2) of human macroscale cortical organization. Raincloud plots show the distributions of G1 and G2 values across *Homo sapiens*-specific and shared cortical regions.

**Figure 5.**
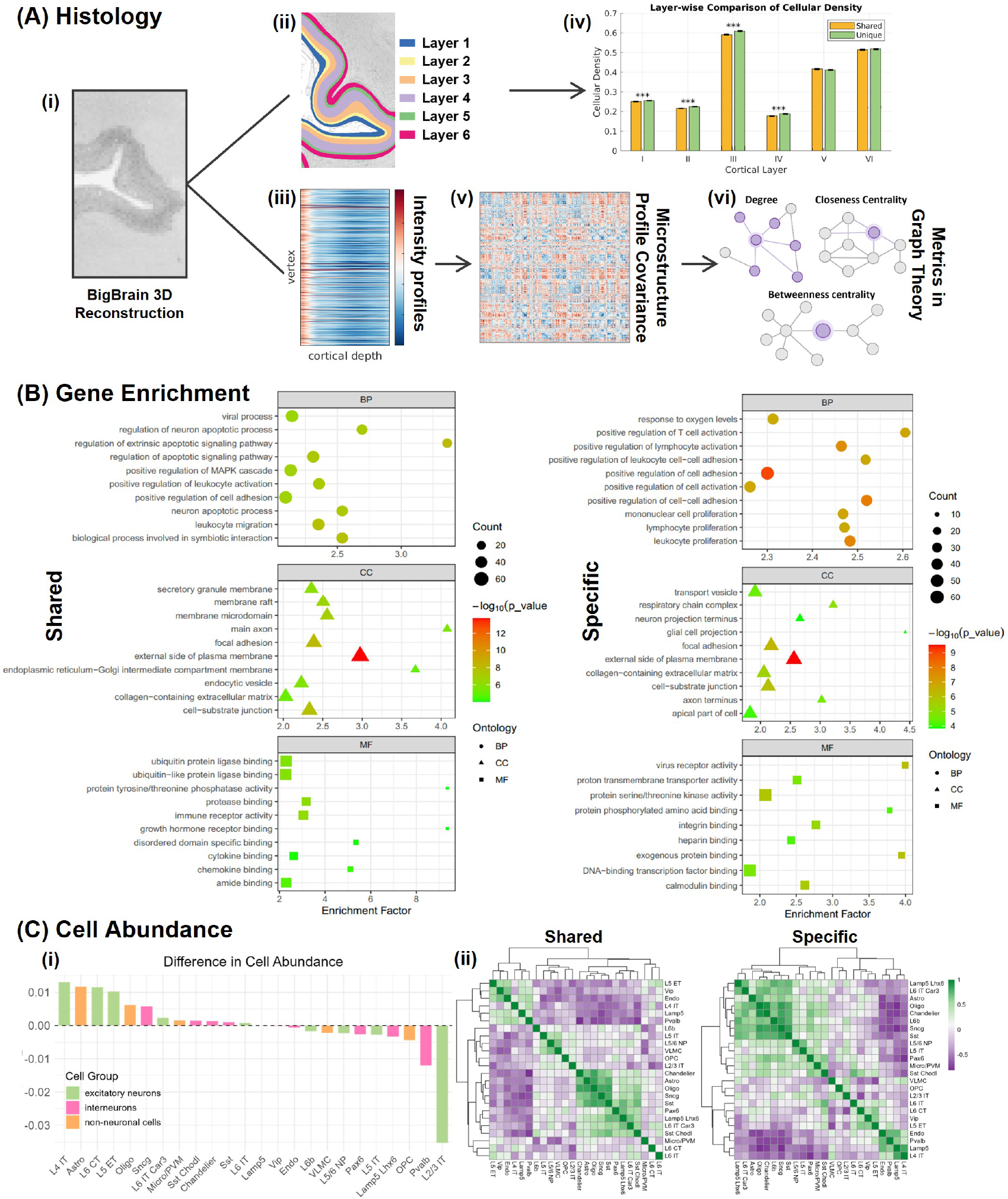
Microstructural differences between *Homo sapiens* and related species revealed by sulcal pit distribution. Histological analyses. (i) BigBrain 3D histological reconstruction; (ii) cortical laminar organization spanning layers I–VI; (iii) sampling of 50 intensity profiles across cortical depth; (iv) comparison of neuronal cell density across six layers between the two cortical region types; (v) construction of microstructural profile covariance (MPC) matrices; and (vi) graph-theoretical analysis of MPC. (B) Genetic analyses. Gene ontology enrichment results for the top 2,000 genes associated with *Homo sapiens*-specific and shared regions. (C) Cellular analyses. (i) Quantification of differences in abundance across 24 cell types between *Homo sapiens*-specific and shared regions. (ii) Hierarchical clustering was performed separately to uncover distinct cellular organization patterns in each region type.

Using Neurosynth, we conducted a meta-analysis for the two categories of cortical regions to identify their associated functional domains (**Fig. 4B**(i)). The analysis reveals that the species-shared cortical regions are mainly associated with terms like ‘nerve’, ‘M1’, and ‘sensorimotor cortex’, which correspond to unimodal and motor-related brain regions. In contrast, *Homo sapiens*-specific cortical regions are primarily associated with terms such as ‘depressed’, ‘personality’, ‘spontaneous’, and ‘communication’, which relate to higher-order cognitive and emotional functions. **Fig. 4B**(ii) presents a statistical analysis of shared and *Homo sapiens*- specific sulcal pit distributions across different brain networks based on the Yeo7 atlas. Higher values indicate that the corresponding brain network exhibits greater degree of cross-species similarity (e.g., SMN), whereas lower values reflect stronger *Homo sapiens* specificity in pit distribution (e.g., DMN). **Fig. 4B**(iii) illustrate the first two principal gradients (G1 and G2) of human cortical macrostructural organization, derived from resting-state functional connectivity data (HCP S1200). Gradient 1 (G1) spans a continuum from transmodal regions (yellow) to unimodal sensory regions (blue). Gradient 2 (G2) captures modality segregation, distinguishing somatomotor and auditory cortices (yellow) from visual areas (blue) [34]. The accompanying raincloud plots visualize the distributions of G1 and G2 values across *Homo sapiens*-specific and shared cortical regions. For G1, *Homo sapiens*-specific regions tend to be localized toward the transmodal area, exhibiting a relatively balanced bimodal distribution with a positive median, while shared regions skew toward unimodal area. For G2, although both categories cluster near zero, shared regions are more frequently localized toward the somatomotor end, whereas *Homo sapiens*-specific regions exhibit a subtle shift toward the auditory direction. Statistical comparisons reveal significant differences in both G1 and G2 distributions between *Homo sapiens*-specific and shared regions (p *<* 0.001).

We utilized post-mortem human brain data from the BigBrain repository (https://bigbrain.loris.ca/main.php [35]) to investigate microstructural differences between *Homo sapiens*-specific and shared cortical regions. The dataset comprises ultra-high-resolution, Merker-stained histological reconstruction of a human brain, where staining intensity reflects neuronal density and soma size (**Fig. 5A**(i)). We quantified neuronal density across the six cortical layers within both *Homo sapiens*-specific and shared cortical regions. As shown in the accompanying bar plot, *Homo sapiens*-specific regions exhibit significantly higher neuronal density in the superficial layers (layers I–IV) compared to shared regions. In contrast, no significant differences were observed in the deeper layers (layers V–VI) (**Fig. 5A**(iv)). These findings suggest that microstructural divergence between human-specific and conserved cortical areas is primarily concentrated in the upper cortical layers. To investigate microstructural connectivity differences between the two types of cortical regions, we sampled cortical cell-staining intensity profiles of 50 equidistant layers (**Fig. 5A**(iii)) from the BigBrain dataset as a feature and employed the schaefer-100 atlas to define cortical nodes, thereby establishing a microstructural profile covariance (MPC) (**Fig. 5A**(v), details see Methods). We computed graph-theoretical metrics based on the MPC, including measures of degree, local clustering, path-based integration, and community structure. Across all metrics, *Homo sapiens*-specific regions showed higher values than species-shared regions. This suggests that *homo*-specific cortical regions are characterized by greater local integration and denser microstructural connectivity, potentially supporting more complex functional integration.

We utilized cortical gene expression data from the Allen Human Brain Atlas (AHBA) to examine the expression differences between shared and *Homo*-specific cortical regions. Using partial least squares regression (PLSR), we identified the top 2,000 genes most strongly associated with each region type, followed by Gene Ontology (GO) and Kyoto Encyclopedia of Genes and Genomes (KEGG) enrichment analyses (**Fig. 5B** and Fig.S8). In the enrichment plot, the x-axis represents the enrichment factor, color denotes the significance level (p value), and dot size indicates the number of genes within each GO term. In species-shared cortical regions, enriched GO terms such as regulation of neuron apoptotic process, positive regulation of cell adhesion, regulation of apoptotic signaling pathway, and leukocyte migration indicate strong conservation of fundamental processes related to structural maintenance and immune surveillance. By contrast, *Homo*-specific regions are enriched for positive regulation of T cell activation, response to oxygen levels, and mononuclear cell proliferation, reflecting a shift toward immune modulation and metabolic adaptation. These patterns indicate that *Homo*-specific cortical areas may have undergone evolutionary adaptation toward integrating internal physiological states (e.g., immune and oxygen signals) with brain activity, potentially supporting complex socio-emotional and self-regulatory functions unique to *Homo sapiens*. KEGG pathway analysis revealed genes in species-shared regions were mainly enriched in pathways related to neurodegenerative and metabolic processes, involving fundamental neuronal functions such as mitochondrial activity, oxidative phosphorylation, ROS metabolism, and apoptosis regulation. This pattern indicates that shared regions retain conserved mechanisms of neuronal energy metabolism across species. In contrast, *Homo*-specific regions were enriched in signaling and synaptic pathways, including dopaminergic and glutamatergic transmission and MAPK signaling, suggesting enhanced molecular specialization for neural regulation and higher cognitive functions in humans.

To delve deeper into the microstructural differences between the two regions, we analyzed the variations in cell abundances. Brain cells can be categorized into three classes and 24 subclasses (details is provided in the Supplementary Information). We quantified the abundance differences of each cell type between specific and shared regions (**Fig. 5C(i)**. Green colors represent excitatory neurons, pink colors represent interneurons, and orange represent non-neuronal cells. In terms of cell-type abundance differences, excitatory neurons show a more pronounced bimodal distribution between specific and shared regions. In contrast, interneurons and non-neuronal cells display relatively modest differences between the two region types. Among excitatory neurons, specific regions exhibit higher abundances of deep-layer projection neurons such as L4 IT, L5 ET, and L6 CT, suggesting enhanced capacity for integrative processing and long-range communication to support complex cognitive functions. Conversely, shared regions show higher proportions of L2/3 IT neurons, which are more involved in local circuit processing and modular network organization. For interneurons, shared regions tend to rely more on fast-spiking synaptic inhibitors such as Pvalb cells, maintaining precise and stable information transmission. In contrast, specific regions are enriched with modulatory interneurons such as Sst and Vip, which may support fine-tuned, flexible regulation of neural activity during higher-order cognitive processing.

To investigate the internal consistency and heterogeneity of cellular composition across cortical regions, we constructed similarity matrices based on the abundance profiles of 24 cell types in *homo*-specific regions and species shared regions, and visualized their clustering patterns using hierarchical clustering (**Fig. 5C**(ii)). The shared region exhibited a more modular and internally coherent structure, where distinct cell clusters were readily observed. For instance, some excitatory neurons (such as L5 ET, L4 IT) and interneurons (such as Vip, Endo) were grouped into a tight cluster, while non-neuronal cells like Astro and Oligo formed another clear module. This indicates a stable and conserved cellular organization across shared cortical areas. In contrast, the specific region showed a more complex and heterogeneous clustering structure, with broader correlation distributions and less distinct module boundaries. The appearance of both excitatory cells (e.g., L6b, L6 IT Car3) and inhibitory cells (e.g., Lamp5 Lhx6, Chandelier) and non-neuronal (e.g. Astro, Oligo) within mixed clusters supports the notion that specific regions may adopt more specialized or adaptive cellular arrangements to support complex *homo*-specific cognitive functions. These findings suggest that shared regions maintain a conserved and modular cell type architecture, while *homo*-specific regions exhibit higher heterogeneity and functional integration, potentially reflecting evolutionary adaptations.

## Discussion

Understanding brain evolution requires approaches that are both biologically meaningful and comparable across species. Traditional atlas-based metrics are limited by species-specific parcellations, reducing their cross-species applicability. Sulcal pits, as high-resolution and atlas-independent cortical landmarks, provide a robust atlas-independent alternative that can be consistently applied across both extant and extinct species. Using cortical data from 90 mammals, we built a first cross-species sulcal pit database, revealing patterns linked to phylogeny and ecology. Evolutionary analyses identified primate–non-primate divergence and species-specific morphological inflection points. In humans, *Homo*-specific cortical regions are associated with higher cognition and emotion, and show unique microstructural, genetic, and cellular traits, highlighting sulcal pits’ value in bridging cortical morphology and evolutionary neuroscience.

Sulcal pits correspond to the locally deepest points within cortical sulci, and their number and spatial arrangement collectively capture the feature of cortical folding. At the most fundamental level, whole-brain pit count provides an intuitive yet quantitative measure of folding complexity. Among primates, this complexity spans a broad range—from the highly convoluted *Homo sapiens* cortex, which contains the greatest number of pits (57 in the left hemisphere), to the much simpler brain of the prosimian *Galago senegalensis*, which possesses only three. In contrast, rodents exhibit markedly fewer pits overall and minimal interspecies variation, consistent with their predominantly lissencephalic cortical architecture. We also found significant variations between social and non-social species, as well as fossorial and non-fossorial species. Social species tend to exhibit higher pit counts compared to non-social counterparts, and non-fossorial species generally have more pits than fossorial species. Our analysis revealed systematic variation in sulcal pit counts across taxonomic and ecological dimensions. Primates exhibited more pits than rodents, consistent with their generally higher cortical folding complexity and expanded surface area, features linked to enhanced functional specialization and cognitive capacity [7]. Species living in large social groups also showed higher pit counts than solitary or pair-living species, in line with the social brain hypothesis that complex social interactions select for increased cortical differentiation [36]. Conversely, fossorial species displayed markedly fewer pits than non-fossorial species, suggesting that life in constrained, low-visibility environments may reduce demands for sensory integration and flexible cognition. These patterns indicate that the whole-brain pit count capture both phylogenetic heritage and ecological adaptation, providing an evolutionarily informative marker of cortical organization [37]. Cortical surface area and brain volume are commonly used indices in comparative neuroanatomy to represent brain size. Although pit counts showed positive allometry with both measures, a substantial proportion of variance (∼ 20%) remained unexplained. After controlling for surface area and volume, significant differences in size-adjusted pit counts persisted across taxonomic and ecological categories. These findings indicate that sulcal pit count is not solely determined by brain size but reflects additional lineage- and ecology-specific factors, providing complementary information beyond traditional size-based metrics.

The sulcal pit distribution similarity matrix exhibited a significant positive correlation with the phylogenetic similarity matrix (Mantel test: *r* = 0.35, *p* = 0.001), indicating that species with closer evolutionary relationships tend to exhibit more similar pit organization patterns. This association was stronger in primates (*r* = 0.41, *p* = 0.001) than in rodents (*r* = 0.11, *p* = 0.04), despite primates displaying greater inter-species variation in cortical folding complexity. These findings demonstrate that sulcal pits, as early-forming landmarks of cortical folding, are subject to strong phylogenetic constraints, with their spatial distribution preserving signatures of evolutionary history across species [29]. A likely explanation lies in the developmental origin of sulcal pits: their formation is closely linked to neuronal proliferation and migration during early cortical development—processes strongly regulated by genetic factors—thus allowing their macroscale organization to remain relatively stable over evolutionary timescales [38, 28]. Although primates exhibit greater interspecies variation in cortical folding complexity compared to non-primates (Fig.2A, right), the correlation between pit distribution similarity and phylogenetic relatedness is notably stronger in primates (Fig.2A, left). One possible explanation is that major morphological divergences among primates occurred relatively recently, around 20 million years ago, whereas in non-primates—particularly rodents—such evolutionary shifts were concentrated around 30 million years ago [39, 40]. The more recent divergence in primates may have preserved a greater proportion of shared early folding patterns, thereby strengthening the phylogenetic signal in pit distribution [41]. Another contributing factor may be ecological diversity: rodents occupy a wider range of habitats and experience more heterogeneous environmental pressures, which could promote greater divergence in pit organization even among closely related species, thereby weakening the association between pit similarity and phylogenetic proximity [42, 40]. Taken together, these results indicate that sulcal pit architecture encodes lineage-specific constraints that are more strongly preserved in primates, highlighting its potential as a stable morphological marker for evolutionary neuroscience.

Principal component analysis of the pit distribution similarity matrix further revealed that species were organized along low-dimensional axes corresponding closely to their phylogenetic and ecological affiliations. The first two components explained 33.4% and 15.6% of the total variance, and the primary axis (PC1) largely separated primates from non-primates. This result suggests that sulcal pit patterns embed phylogenetic information in a compact morphometric space, highlighting the capacity of pit-based representations to recapitulate evolutionary structure without requiring atlas alignment. To test whether the correspondence between pit similarity and phylogenetic relatedness could be explained by differences in brain size, we applied multiple regression on distance matrices (MRM) [31], which simultaneously evaluated the effects of several predictors on pairwise dissimilarity (Fig. 2C). The model included phylogenetic distance, cortical surface area difference, and brain volume difference as predictors. Consistent with the Mantel results, phylogenetic distance remained the only significant predictor of pit dissimilarity, whereas cortical surface area and brain volume differences were not significant. These findings confirm that the observed pit–phylogeny correspondence is robust and independent of allometric scaling effects, reinforcing that pit architecture captures lineage-specific organizational principles rather than mere consequences of brain enlargement. Finally, hierarchical clustering based on pit distribution similarity produced a species arrangement that approximated the phylogenetic tree, with primates and non-primates mostly grouped into distinct clusters even without explicit phylogenetic constraints. This demonstrates that the spatial organization of sulcal pits inherently encodes phylogenetic structure.

In the evolutionary trajectories of cortical folding, primates typically exhibit a gradual early increase followed by rapid accumulation, with evolutionary turning points concentrated around 20 Mya. In contrast, rodents more often display an initial decline followed by stabilization, with earlier turning points (∼ 30 Mya) that are more widely distributed. This divergence may reflect distinct strategies in ecological niche occupation, metabolic demands, and the evolution of higher cognitive abilities across clades. In primates, prolonged neurogenesis, enlarged progenitor pools, and enhanced radial and tangential neuronal migration provide the developmental substrate for progressive cortical elaboration and increased gyrification [38, 43, 44]. These biological mechanisms enable the formation of new association areas and hierarchical cortical networks, supporting advanced functions such as social cognition, fine motor control, and visual integration—features that drive the accelerated accumulation of folding complexity during later evolutionary phases. In rodents, early niche diversification was accompanied in some lineages by a phenomenon of ‘secondary lissencephaly’, wherein previously folded brains evolved toward a smoother phenotype [43]. This trend may be associated with secondary loss of miR-3607 [45], which reduces neuronal numbers and decreases cortical complexity [46], potentially representing an energy-optimizing strategy for specific ecological contexts [47]. Because sulcal pit formation is established during early cortical development and strongly regulated by genetic factors, species that are phylogenetically closer tend to exhibit greater similarity in cortical complexity [48] as well as in the timing of evolutionary turning points. Our findings support this view, showing that differences in turning point timing across species are positively correlated with phylogenetic distance, with particularly close correspondence within primates. This suggests that primates have retained a stronger phylogenetic signal in their evolutionary trajectories, a pattern also observed in other taxa [49]. In terms of branching evolutionary patterns, primates generally show increased cortical complexity from ancestor to descendant lineages, whereas non-primates do not consistently follow this trend. This may be due to increased neuronal density following primate divergence, driven by advanced cognitive demands, which in turn enhances cortical folding complexity at the macroscale [44]. In contrast, some non-primate species may simplify folding structures under certain environmental pressures to reduce metabolic costs (e.g., in small rodents). Moreover, the broad geographic range and long migratory distances typical of rodents may favor reduced cognitive investment to balance energy budgets, a pattern analogous to brain size reductions seen in migratory versus resident birds [50].

At the functional network level, primates exhibit the fastest evolution in the Ventral Attention Network (VAN) and the Default Mode Network (DMN), likely reflecting high demands on attentional reorienting and self-referential processing required by complex social interactions and environmental exploration [51, 52]. Non-primates, by contrast, show the most rapid evolution in the Limbic Network, which plays a central role in innate social, emotional, and motivational behaviors critical for survival [53]. The DMN evolved rapidly in both clades [51], consistent with evidence that it is both conserved within species and subject to unique, rapid adaptive changes [54], potentially reflecting its universal role in multimodal integration and internal modeling. Finally, regarding the coupling between surface area expansion and folding complexity, primates generally display strong positive correlations, especially in the DMN, where expansion is almost invariably accompanied by substantial increases in folding complexity to support greater connectivity and information integration. In contrast, non-primates show weaker coupling overall, with the VAN even exhibiting a negative correlation. This may be because, in non-primates, VAN (often overlapping with parts of the salience network, including the anterior insula and ventrolateral prefrontal cortex) relies more on midbrain and brainstem circuits for rapid, low-latency sensory integration (e.g., thalamocortical and corticobasolateral amygdala pathways), rather than on increased cortical folding to enhance local processing capacity. Such a strategy may favor reduced folding as surface area increases, shortening conduction pathways and resulting in a negative correlation.

Previous analyses demonstrated that sulcal pits can reveal pronounced differences in brain evolution. However, such evidence alone is insufficient to establish pits as a universally applicable tool for evolutionary studies. To more rigorously test their capacity to detect fine-grained evolutionary differences, we narrowed the scope from broad taxonomic comparisons to closely related species pairs, selecting six primate species with the smallest phylogenetic distance to *Homo sapiens*. We then performed multi-scale analyses spanning functional, histological, transcriptomic, and cellular levels to investigate the structural and functional features of *Homo*-specific cortical regions. Overall, our findings show that *Homo*-specific regions are predominantly located in higher-order transmodal cortices, particularly within the default mode network (DMN) and ventral attention network (VAN), which are associated with self-referential processing, emotional regulation, language, and social interaction. In contrast, species shared regions are concentrated in unimodal sensory and motor cortices. This spatial distribution aligns with the view that advanced cognitive functions depend on the integration of information across multiple modalities [55, 34, 56], suggesting that sulcal pits encode not only morphological variation but also evolutionary signals of cognitive specialization.

At the histological level, BigBrain data reveal that *Homo*-specific regions have significantly higher neuronal density in the superficial cortical layers (I–IV) compared to shared regions, with no significant differences in the deeper layers (V–VI). Superficial-layer neurons are primarily involved in integrating inputs from different sensory modalities and cortical areas, as well as in cross-domain associative processing, whereas deep-layer neurons mainly transmit cortical outputs to subcortical structures or distant cortical regions [57, 58, 59]. The higher density of superficial neurons in *Homo sapiens*-specific regions suggests a stronger microstructural basis for multimodal integration and higher-order cognitive processing [60]. Graph-theoretical analyses of microstructural profile covariance further indicate that *Homo*-specific regions exhibit greater node degree, closeness centrality and betweenness centrality, reflecting denser local integration and richer interareal connectivity, potentially supporting complex cognitive operations [61, 62]. At the molecular level, gene expression analyses reveal that shared regions are enriched for conserved processes such as cell adhesion, regulation of apoptosis, and immune surveillance [63], whereas *Homo*-specific regions are enriched for pathways related to T cell activation, oxygen-level response, and mononuclear cell proliferation—reflecting immune modulation and metabolic adaptation [64]. This pattern may indicate a co-evolution of brain function and the immune system, in which immune signals not only maintain neural homeostasis but also influence synaptic plasticity and network activity, thereby modulating emotion and social behavior [65]. Such enrichment in immune modulation and metabolic adaptation pathways may provide a molecular basis for the human-specific cortical regions to support complex socio-cognitive functions [66]. At the cellular level, *Homo*-specific regions show a higher proportion of deep-layer projection excitatory neurons (L4 IT, L5 ET, L6 CT), supporting long-range information transmission and integration across multiple brain regions [67, 68], whereas shared regions contain a higher proportion of superficial, locally processing L2/3 IT neurons, favoring modular sensorimotor processing—consistent with the findings from our histology-based analyses [69]. In terms of cellular organization, hierarchical clustering results show that shared regions exhibit a clear modular structure and high internal consistency, whereas *Homo*-specific regions have blurred boundaries, greater heterogeneity, and mixed distributions of multiple cell types. This ‘demodularized’ pattern may provide a cellular basis for cross-domain information integration and multitasking. Previous studies have shown that the expanded association cortex in human contains a more diverse repertoire of cell types, which are key hubs supporting higher-order cognitive functions. Brain regions with more complex functional demands tend to harbor a richer diversity of neuronal subtypes to enable more flexible computational capacitie [70], consistent with the high heterogeneity observed in our study.

## Methods

### Dataset description

#### Cortical Surfaces Across Species

In this study, we utilized publicly available cortical surface data from 90 mammalian species along with their corresponding ecological data, including 58 primates, 28 rodents, 2 lagomorphs, 1 scandentian, and 1 dermopteran, as provided by [71]. The original data combines imaging from various sources and modes, with 75 specimens imaged via MRI, 7 via diffusion iodine contrast-enhanced computed tomography (DiceCT), and 8 through serial histological sections. Ecological and behavioral data were sourced from the literature, covering aspects such as diurnal activity patterns, social group size, and preferred habitat. Arboreal-spends most of its time in trees, in a cluttered environment, is able to move in all directions; Terrestrial-spends most of its time on the ground, in an open-environment, does not habitually move in vertical direction; Fossorial-spends most of its time underground, affecting its range of movement and perception. In cases of ambiguity in the literature, species were assigned to multiple ecological categories. For activity patterns, species described as ‘diurnal’, ‘cathemeral’, or ‘arrhythmic’ were uniformly classified as ‘diurnal’. Social structure was categorized as ‘large social group’ for species not explicitly described as solitary or pair-living.

To achieve cortical surface alignment across species, Schwartz [71] constructed a cross-species common reference frame for cortical geometry through a systematic pipeline. First, cortical surface models were derived via manual segmentation of imaging data from 90 species. Pairwise correspondences between sister species’ cortical surfaces were then established using computational geometry methods: intrinsic spectral features were integrated with extrinsic morphological descriptors to encode local cortical shape, and dense alignments were achieved by matching these multi-modal features in a high-dimensional embedding space via Coherent Point Drift (CPD). Ancestral cortical shapes were reconstructed via shell-space interpolation, iteratively applied along the phylogenetic tree up to the common ancestor. All surfaces were subsequently resampled to a unified topology (fsaverage6) to ensure global cross-species correspondence. Surface models are available at https://github.com/cirmuw/EvolutionOfCorticalShape.

#### Gene Expression Data

The Allen Human Brain Atlas (AHBA) microarray dataset consists of 3,702 spatially distinct tissue samples obtained from six neurotypical adult human brains. In the AHBA, each gene probe is assigned a numerical identifier and a platform-specific label. When a probe uniquely represents a gene, it is further annotated with gene-specific information such as gene symbols and Entrez Gene IDs—stable identifiers curated by the Entrez Gene database at the National Center for Biotechnology Information (NCBI). This probe-level dataset provides high-resolution transcriptional coverage across nearly the entire brain, offering expression measurements for over 20,000 genes. Preprocessing of the AHBA dataset was conducted following the standardized pipeline detailed in [72], resulting in a region *x* gene expression matrix suitable for downstream regional analyses. Since complete right hemisphere data were available from only two donors, we restricted our analyses to the left hemisphere to ensure consistency and data quality.

#### Cortical snRNA-seq data

We used cell-type abundance estimates provided by Jorstad dataset [73], comprising 24 cortical cell types. These cells include nine GABAergic inhibitory interneurons (PAX6, SNCG, VIP, LAMP5, LAMP5 LHX6, Chandelier, PVALB, SST CHODL and SST), nine glutamatergic excitatory neurons (L2/3 IT, L4 IT, L5 IT, L6 IT, L5 ET, L5/6 NP, L6 CT, L6b and L6 IT Car3), and six non-neuronal cells (Astro, Endo, VLMC, Oligo, OPC and Micro/PVM). The abundances were represented in fsaverage6 surface space and subsequently parcellated into the Schaefer 100 atlas. For each cortical parcel, values were averaged across vertices within individual donors and then across donors to yield parcel-level cell-type abundance profiles.

#### BigBrain

The BigBrain is an ultra-high-resolution 3D volumetric histological reconstruction of a postmortem human brain (https://bigbrain-ftp.loris.ca/). The brain was paraffin-embedded, sectioned coronally into 20-µm slices, silver-stained for cell bodies, and digitised. Artefacts such as rips and stain crystallisation were manually inspected and repaired using automated procedures, including alignment to postmortem MRI, intensity normalisation, and block averaging [35]. The reconstructed 3D volume was generated using a coarse-to-fine hierarchical approach and is available at multiple resolutions (100-, 200-, 300-, and 400-µm isovoxel sizes). To quantify cytoarchitectonic features, we analyzed inverted images, where staining intensity reflects cellular density and soma size. The six cortical layers of BigBrain had previously been segmented using a convolutional neural network, and surface reconstructions of the laminar boundaries were publicly available [74].

### Sulcal pits extraction

Based on prior research, sulcal pits were defined as the deepest points of sulci on the cortical surface. For each species, we first computed the 3D convex hull of the cortical mesh using ConvexHull [75]. To visually illustrate the spatial relationship between the cortex and its convex hull, we selected a representative crosssectional view using *Homo sapiens* and *Ochotona macrotis* as examples (Fig.1A). The sulcal depth of each vertex on the cortical surface was calculated as the shortest Euclidean distance to the convex hull surface, visualized by a color gradient in Fig.1B. Local maxima of sulcal depth were identified as sulcal pits and marked by yellow spheres.

To enable cross-species comparison, all cortical surfaces were resampled to the fsaverage6 template (40,962 vertices). Since sulcal pits are defined on the neocortical surface, and in some species the olfactory bulb is anatomically separate from the cortical sheet, we removed the olfactory bulb from those species to ensure accurate convex hull computation. A uniform search radius, 4-ring neighborhood on the surface mesh, was applied to all species to ensure methodological consistency [30]). Since each species was represented by a single cortical surface, intra-species variability was not the focus of our analysis and thus considered negligible. Since the original surfaces correspond to the gray matter boundary, sulcal pits locations are not directly visible. To visualization purposes, *homo* sulcal pits were projected onto the fsaverage6 surface of human (Fig.1B).

### Allometric scaling of sulcal pit count with brain size

To assess the allometric relationship between whole-brain pit count and brain size (cortical surface area, brain volume), we plotted log_10_(pit count) against log_10_(area) and log_10_(volume) with linear fits, colorcoding points by order (primata and non-primate). To control for brain-size scaling, we fit a multivariable log-linear model:

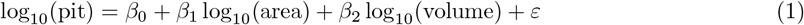

Residuals from this model were defined as the residual pit count:

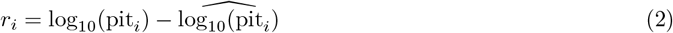

which quantifies each species’ deviation from the expected pit number given its brain size. For each binary factor (order, sociality, fossoriality, terrestriality, arboreality), we compared residuals between the two levels using two-sided Wilcoxon rank-sum tests. After controlling for cortical surface area and brain volume, residual differences between groups (e.g., primata vs non-primate; arboreal vs non-arboreal) remained significant, indicating that pit count carries order specific information independent of overall brain size.

### Phylogenetic similarity matrix (PS)

To quantify evolutionary relationships among species, we constructed a phylogenetic similarity matrix based on the established species phylogeny. Specifically, a time-calibrated phylogenetic tree was obtained from [71], which encodes the evolutionary divergence times between species. For each species pair, we traced their lineages to identify the most recent common ancestor (MRCA) and computed the patristic distance as the sum of the branch lengths from each species to the MRCA, the illustration of the phylogenetic distance between two example species is provided in Supplementary Information Fig.S3. The resulting 90 *×* 90 distance matrix was then reordered according to the leaf order of the phylogenetic tree and converted into a similarity matrix by subtracting each entry from the maximum distance and linearly normalizing the values to the range [-1,1]. In this similarity matrix, larger values indicate closer evolutionary proximity, effectively capturing the degree of phylogenetic relatedness.

When assessing differences between *Homo sapiens* and its closest relatives, we filtered species by phylogenetic distance and identified six species with evolutionary tree distances to *Homo sapiens* of less than 26 million years, meaning that the earliest split between *Homo sapiens* and any of these species occurred no earlier than 13 million years ago. The species names and their distances to *Homo sapiens* are listed in the Supplementary Information.

### Pit distribution similarity matrix (PDS)

For each pair of species, we iteratively examined every sulcal pit in each species to identify the nearest sulcal pit on the cortical surface mesh of the other species and recorded their geodesic distance. The mean of the matching proportions in both directions was then computed as the similarity index of sulcal pit distributions between the species pair. Diagonal elements were set to 1 to indicate perfect self-similarity within the same species.

### Multiple regression on distance matrices

To test whether the association between pit distribution similarity and phylogenetic similarity could be explained by overall brain size or cortical surface area, we employed multiple regression on distance matrices (MRM) [31]. In this framework, each pair of species (*i, j*) contributes one observation, where the dependent variable *Y*_*ij*_ represents the pit dissimilarity between species *i* and *j*, defined as 1 − *r*_*ij*_ with *r*_*ij*_ being the Pearson correlation of pit maps. The predictors included phylogenetic distance ( 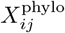, derived from patristic distances on the evolutionary tree), cortical surface area difference ( 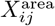, Euclidean distance), and brain volume difference ( 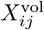, Euclidean distance). The model can be expressed as:

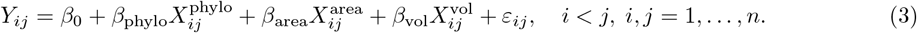

Because dyadic observations are not independent, significance was assessed using a quadratic assignment procedure (QAP), which simultaneously permutes rows and columns of *Y*_*ij*_ to generate a null distribution of regression coefficients. We performed 10,000 permutations for each model. The full results for each predictor, including standardized *β*, permutation *p* values, raw *β*, and corresponding *p* values, are summarized in Supplementary Table S1.

### Hierarchical clustering

To compare phylogenetic ordering with an unsupervised organization derived from pit-based similarity, we first computed a species-by-species pit distribution similarity matrix *PDS* ∈ ℝ^90*×*90^ (see *Pit similarity* for details). For clustering, *S* was converted to a dissimilarity matrix *D* = **1**− *PDS*, where larger values indicate lower similarity. We then applied agglomerative hierarchical clustering with average linkage (UPGMA) to *D* and obtained a one-dimensional species ordering using optimal leaf ordering to maximize adjacent similarity. No phylogenetic information was used in this procedure. For visualization, the top panel of Fig. 2C displays species ordered by the evolutionary tree, and the bottom panel displays the ordering returned by hierarchical clustering of *D*. The same set of species and annotation bars (taxonomic order, group living, habitat) were used for both panels to enable direct visual comparison.

### Piecewise linear regression model

We used a piecewise linear regression model to characterize the temporal trajectory of the total number of sulcal pits across the whole brain for each species from 80 Mya to the present. In this model, time (*x*) was treated as the independent variable and the total sulcal pit count (*y*) as the dependent variable. For a single breakpoint *c*, the model is expressed as:

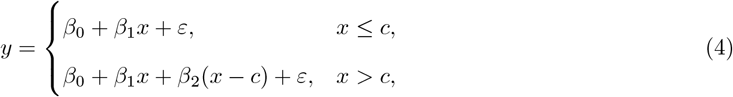

where *β*_0_ is the intercept, *β*_1_ is the slope before the breakpoint, *β*_1_ + *β*_2_ is the slope after the breakpoint, *c* is the breakpoint location (estimated from the data), and *ε* is the error term. The breakpoint *c* was interpreted as the evolutionary turning-point time in brain complexity, marking a change in the rate of sulcal pit evolution. This formulation allows changes in slope at specific evolutionary time points, enabling us to detect stage-dependent patterns in the evolution of sulcal pit counts.

### Generalized additive model

We applied generalized additive model (GAM) to characterize the temporal trajectories of total sulcal pit counts for primate and non-primate groups from 80 Mya to the present. For each group, pit counts along phylogenetic lineages were normalized to a [0, 1] range and fitted using a GAM with time as the predictor and a smooth term *s*(time, *k* = 3) to allow for low-complexity nonlinear trends. Smoothed evolutionary curves were derived from the fitted models, and model fit was summarized by the percentage of deviance explained (pseudo-*R*^2^).

### Surface area expansion rate

The cortical surfaces of 90 species were parcellated into seven functional regions using the Yeo7 template. For each region, the total area of its triangular faces was computed and normalized by the total cortical surface area. Using the common ancestor as a reference, the surface area expansion rate for each region was defined as the difference between the area of that region in a given species and the corresponding area in the ancestor.

### *Homo sapiens*-specific and species-shared sulcal pits

To identify *Homo sapiens*-specific and species-shared sulcal pits, we first aggregated the pits from six closely related species onto a common standardized cortical template, creating the sulcal pit pattern of *Homo sapiens*’ closest relatives. These six species were selected based on their evolutionary distance to *Homo sapiens*, as determined by pairwise phylogenetic distances. We then iterated through all pit vertices on the standardized cortical surface of *Homo sapiens*. For each pit, a 5-ring neighborhood on the cortical mesh was used as the search radius to determine the number of pits from the reference pattern occurring within this region. The pit was then assigned a shared label equal to this count, with the resulting shared label ranging from 1 to 3, or assigned a label of –1 if no reference pits were detected. The resulting distribution of labels is shown in Fig. **4A**(ii).

### Meta-analysis on cognitive functions

To identify the functional correlates of the *Homo sapiens*-unique and species-shared cortical regions, we performed meta-analytic decoding using Neurosynth (http://www.neurosynth.org) [76]. Neurosynth integrates large-scale meta-analysis with text mining to infer probabilistic associations between cognitive terms and spatial activation patterns. We applied this analysis to the cortical regions that are unique to *Homo sapiens* and those shared with closely related species, respectively. For each region type, we visualized the top cognitive terms most strongly associated with their spatial distribution.

### Functional gradients

To illustrate how *Homo sapiens*-unique and species-shared regions are involved in brain functional organization, we referenced large-scale macroscale functional gradients in the human brain. Specifically, we utilized the group-level functional connectivity (FC) matrix derived from the Human Connectome Project (HCP), based on time series extracted from the schaefer-100 parcellation. Pairwise Pearson correlation coefficients were computed to obtain individual FC matrices, which were then averaged across subjects to generate a group-level FC matrix. For each row of the matrix, we retained the top 10% of connections and computed the row-wise normalized angle similarity. The resulting similarity matrix was then subjected to diffusion map embedding to derive the principal gradient of functional connectivity, capturing the dominant axis of macroscale functional organization [34].

### Histology-based microstructural profile covariance

We constructed 50 equivolumetric intracortical surfaces to sample staining intensities across cortical depths [77]. This yielded vertex-wise intensity profiles reflecting microstructural composition of each cortical vertex. Data sampled from surfaces closest to the pial and white matter boundaries were discarded to mitigate partial volume effects. Vertex-wise intensity profiles were subsequently averaged within 100 functionally defined parcels [78]. Microstructural Profile Covariance (MPC) was computed by calculating Pearson’s correlations between the averaged intensity profiles of each pair of regions.

### Partial least squares regression

In the partial least squares regression (PLSR) framework, the input matrix *X* ∈ ℝ^*n×p*^ represents regional gene expression (*n* = 1000 cortical regions, *p* = 15,631 genes), and the response vector *y* ∈ ℝ^*n*^ encodes evolutionary labels (−1 for *Homo sapiens*-specific regions, and 1–3 for species-shared regions with increasing conservation). Gene expression values were *z*-scored across regions, and missing entries were imputed from the nearest valid neighbor.

The number of components in the PLSR model was determined using 10-fold cross-validation, selecting the value that minimized the mean squared error (MSE). For both *Homo sapiens*-unique and species-shared regions, the MSE reached its minimum at component = 10, which was therefore chosen for the final model (see Supplementary Information, Fig.S7 for the cross-validation curves). PLSR was then performed as:

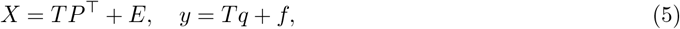

where *T* ∈ ℝ^*n×h*^ denotes latent scores, *P* ∈ ℝ^*p×h*^ loadings for genes, and *q* ∈ ℝ^*h*^ loadings for the response. Gene contributions were computed as the weighted sum of absolute loadings across components, with weights proportional to the variance explained by each component. Genes were ranked by contribution, and the top 2000 genes were retained separately for *Homo*-specific and species-shared regions for downstream enrichment analysis.

### Gene enrichment

To explore the potential molecular mechanisms underlying the spatial distribution of *Homo sapiens*-unique and species-shared cortical regions, we applied partial least squares regression to relate interregional gene expression patterns (derived from the Allen Human Brain Atlas [72]) to the spatial maps of these two cortical categories. For each category, we identified the top 2000 genes that showed the highest absolute loadings on the first PLS component, indicating the strongest association with the spatial pattern of interest. These gene sets were then subjected to gene enrichment analysis using the clusterProfiler toolkit[79], targeting three major Gene Ontology (GO) domains: Biological Process (BP): describing the biological objectives to which the genes contribute; Cellular Component (CC): specifying where the gene products are active within the cell; and Molecular Function (MF): characterizing the biochemical activities performed by gene products.

### Statistical treatment

The distribution of variables was inspected for normality using the Shapiro–Wilk test. For variables following a normal distribution and with comparable variances, group comparisons were performed using two-sample Student’s t-tests. To account for multiple comparisons, all p-values were corrected using the false discovery rate (FDR) method, with *q <* 0.05 considered significant. In cases of unequal variances or unbalanced sample sizes, Welch’s t-test was applied. For variables violating the normality assumption, we employed the non-parametric Wilcoxon rank-sum test (Mann–Whitney U test) to evaluate whether two independent samples originated from the same distribution.

For correlation analyses, Pearson’s correlation coefficient (PCC) was computed depending on the distribution of the data. In addition, Mantel tests were used to evaluate the correspondence between similarity/distance matrices (e.g., phylogenetic versus pit distribution similarity). For modeling temporal trajectories, generalized additive models (GAMs) were fitted with evolutionary time as predictor, and model fit was evaluated using the percentage of deviance explained (pseudo-*R*^2^). Hierarchical clustering was performed on dissimilarity matrices derived from pit distribution similarity using agglomerative linkage with optimal leaf ordering. Multiple regression on distance matrices (MRM) was conducted to test whether the pit-phylogeny associations could be explained by brain size or ecological covariates. Effect sizes (e.g., Cohen’s *d* for t-tests and rank-biserial correlation for Wilcoxon tests) were reported alongside test statistics where appropriate. Unless otherwise specified, all tests were two-sided.

#### Software

All analysis was implemented in Matlab R2019b and R 4.4.2. Additional processing was performed using FSL 6.0.4, FreeSurfer 6.0.0 and 7.1.1.

### Data availability

All quantitative data supporting the findings of this study are provided as Supplementary Information to the article. Aligned surface models used to define the proposed common reference frame, as well as ancestral state estimates obtained from it are publicly available at https://github.com/cirmuw/EvolutionOfCorticalShape/tree/main/_surfaces. Task-based function association analyses were based on NeuroSynth [76] http://www.neurosynth.org.

### Code availability

All the scripts and visualization are openly available at a GitHub repository (https://github.com/zsy0728/Species-evolution-based-on-pit). The packages are completely open for use, see documentations: BigBrainWarp (https://bigbrainwarp.readthedocs.io/en/latest/), BrainStat (https://brainstat.readthedocs.io/en/), BrainSpace (https://brainspace.readthedocs.io/en/latest/).

## End Notes

## Acknowledgements

Songyao Zhang was supported by the Young Scientists Fund of the National Natural Science Foundation of China (No. 62403103), the Joint Funds of the National Natural Science Fundation of China (No. U24A20753), and the Fundamental Research Funds for the Central Universities (DUT24RC(3)050). Mingrui Zhuang was supported by the China Postdoctoral Science Foundation (No. 2025M772947). Marco Palombo was supported by the UKRI Future Leaders Fellowship MR/T020296/2 and 1073, as well as the MRC Research Grant MR/031566/1. Xi Jiang was partly supported by the National Natural Science Foundation of China (62276050, 62576077) and Sichuan Science and Technology Program (2024NSFSC0655). Tuo Zhang was supported by the National Natural Science Foundation of China (62476222, 62131009). Hongkai Wang was supported by the Joint Funds of the National Natural Science Fundation of China (No. U24A20753), the general program of National Natural Science Fund of China (No. 81971693), the funding of Liaoning Key Lab of IC & BME System and Dalian Engineering Research Center for Artificial Intelligence in Medical Imaging and Liaoning technology innovation center of hyperpolarized MRI.

## Author Contributions

Conceptualization: S.Z.,T.Z. Methodology: S.Z., M.Z. Original Draft: S.Z. Writing - Review & Editing: X.J., T.Z., M.P., H.W. Visualization: S.Z., Y.T., W,Y. Project administration: S.Z. Funding acquisition: S.Z., H.W. Supervision: H.W.

## Declaration of Interests

The authors declare no competing interests.

## Notes

https://doi.org/10.6084/m9.figshare.30382753

### Competing Interest Statement

The authors have declared no competing interest.

## References

[1] Heuer, K. et al. Evolution of neocortical folding: A phylogenetic comparative analysis of mri from 34 primate species. Cortex 118, 275–291 (2019).

[2] Rilling, J. K. Human and nonhuman primate brains: are they allometrically scaled versions of the same design? Evolutionary Anthropology: Issues, News, and Reviews: Issues, News, and Reviews 15, 65–77 (2006).

[3] Dunbar, R. I. & Shultz, S. Evolution in the social brain. science 317, 1344–1347 (2007).

[4] Krubitzer, L. The magnificent compromise: cortical field evolution in mammals. Neuron 56, 201–208 (2007).

[5] Alldritt, S. et al. Brain charts for the rhesus macaque lifespan. BioRxiv (2024).

[6] Ardesch, D. J. et al. Evolutionary expansion of connectivity between multimodal association areas in the human brain compared with chimpanzees. Proceedings of the National Academy of Sciences 116, 7101–7106 (2019).

[7] Van Essen, D. C. et al. Cerebral cortical folding, parcellation, and connectivity in humans, nonhuman primates, and mice. Proceedings of the National Academy of Sciences 116, 26173–26180 (2019).

[8] Griffa, A. et al. Evidence for increased parallel information transmission in human brain networks compared to macaques and male mice. Nature Communications 14, 8216 (2023).

[9] Huang, S. et al. Transbrain: A computational framework for translating brain-wide phenotypes between humans and mice. bioRxiv 2025–01 (2025).

[10] Xia, X., Zeng, X., Gao, F. & Yuan, Z. Mapping cross-species connectome atlas of human and macaque striatum. Cerebral Cortex 33, 7518–7530 (2023).

[11] Chen, A. et al. Single-cell spatial transcriptome reveals cell-type organization in the macaque cortex. Cell 186, 3726–3743 (2023).

[12] Markov, N. T. et al. A weighted and directed interareal connectivity matrix for macaque cerebral cortex. Cerebral cortex 24, 17–36 (2014).

[13] Lein, E. S. et al. Genome-wide atlas of gene expression in the adult mouse brain. Nature 445, 168–176 (2007).

[14] Wang, Y. et al. The chimpanzee brainnetome atlas reveals distinct connectivity and gene expression profiles relative to humans. The Innovation 6 (2025).

[15] Oegema, R. et al. International consensus recommendations on the diagnostic work-up for malformations of cortical development. Nature Reviews Neurology 16, 618–635 (2020).

[16] Pang, J. C. et al. Geometric constraints on human brain function. Nature 618, 566–574 (2023).

[17] Wang, Y., Necus, J., Kaiser, M. & Mota, B. Universality in human cortical folding in health and disease. Proceedings of the National Academy of Sciences 113, 12820–12825 (2016).

[18] Yan, X. The role of cortical midline structure in diagnoses and neuromodulation for major depressive disorder. Psychoradiology 4, kkae001 (2024).

[19] Del-Valle-Anton, L. & Borrell, V. Folding brains: from development to disease modeling. Physiological Reviews 102, 511–550 (2022).

[20] Yin, S. et al. Morphogenesis and morphometry of brain folding patterns across species. bioRxiv 2025–03 (2025).

[21] Ha, T. H. et al. Fractal dimension of cerebral cortical surface in schizophrenia and obsessive–compulsive disorder. Neuroscience letters 384, 172–176 (2005).

[22] Li, G. et al. Mapping longitudinal development of local cortical gyrification in infants from birth to 2 years of age. Journal of Neuroscience 34, 4228–4238 (2014).

[23] Luders, E. et al. A curvature-based approach to estimate local gyrification on the cortical surface. Neuroimage 29, 1224–1230 (2006).

[24] Kroenke, C. D. & Bayly, P. V. How forces fold the cerebral cortex. Journal of Neuroscience 38, 767–775 (2018).

[25] Mota, B. & Herculano-Houzel, S. Cortical folding scales universally with surface area and thickness, not number of neurons. Science 349, 74–77 (2015).

[26] Krubitzer, L. A. & Seelke, A. M. Cortical evolution in mammals: the bane and beauty of phenotypic variability. Proceedings of the National Academy of Sciences 109, 10647–10654 (2012).

[27] Lohmann, G., Von Cramon, D. Y. & Colchester, A. C. Deep sulcal landmarks provide an organizing framework for human cortical folding. Cerebral Cortex 18, 1415–1420 (2008).

[28] Im, K. & Grant, P. E. Sulcal pits and patterns in developing human brains. NeuroImage 185, 881–890 (2019).

[29] Meng, Y., Li, G., Lin, W., Gilmore, J. H. & Shen, D. Spatial distribution and longitudinal development of deep cortical sulcal landmarks in infants. Neuroimage 100, 206–218 (2014).

[30] Zhang, S. et al. Species-shared and-unique gyral peaks on human and macaque brains. elife 12, RP90182 (2024).

[31] McArtor, D. B., Lubke, G. H. & Bergeman, C. Extending multivariate distance matrix regression with an effect size measure and the asymptotic null distribution of the test statistic. Psychometrika 82, 1052–1077 (2017).

[32] Yeo, B. T. et al. The organization of the human cerebral cortex estimated by intrinsic functional connectivity. Journal of neurophysiology (2011).

[33] Vickery, S. et al. The uniqueness of human vulnerability to brain aging in great ape evolution. Science Advances 10, eado2733 (2024).

[34] Margulies, D. S. et al. Situating the default-mode network along a principal gradient of macroscale cortical organization. Proceedings of the National Academy of Sciences 113, 12574–12579 (2016).

[35] Amunts, K. et al. Bigbrain: an ultrahigh-resolution 3d human brain model. science 340, 1472–1475 (2013).

[36] Dunbar, R. I. The social brain hypothesis. Evolutionary Anthropology: Issues, News, and Reviews: Issues, News, and Reviews 6, 178–190 (1998).

[37] Bertrand, O. C., Püschel, H. P., Schwab, J. A., Silcox, M. T. & Brusatte, S. L. The impact of locomotion on the brain evolution of squirrels and close relatives. Communications biology 4, 460 (2021).

[38] Borrell, V. How cells fold the cerebral cortex. Journal of Neuroscience 38, 776–783 (2018).

[39] Pozzi, L. et al. Primate phylogenetic relationships and divergence dates inferred from complete mitochondrial genomes. Molecular phylogenetics and evolution 75, 165–183 (2014).

[40] Fabre, P.-H., Hautier, L., Dimitrov, D. & P Douzery, E. J. A glimpse on the pattern of rodent diversification: a phylogenetic approach. BMC evolutionary biology 12, 88 (2012).

[41] Sathishkumar, R. N. et al. A morphological comparison of the caudal rami of the superior temporal sulcus in humans, chimpanzees, and other great apes. bioRxiv 2025–07 (2025).

[42] Kay, E. H. & Hoekstra, H. E. Rodents. Current Biology 18, R406–R410 (2008).

[43] Kelava, I., Lewitus, E. & Huttner, W. B. The secondary loss of gyrencephaly as an example of evolutionary phenotypical reversal. Frontiers in neuroanatomy 7, 16 (2013).

[44] Lancaster, M. A. Unraveling mechanisms of human brain evolution. Cell 187, 5838–5857 (2024).

[45] Chinnappa, K. et al. Secondary loss of mir-3607 reduced cortical progenitor amplification during rodent evolution. Science advances 8, eabj4010 (2022).

[46] Churakov, G. et al. Rodent evolution: back to the root. Molecular biology and evolution 27, 1315–1326 (2010).

[47] Aristide, L. et al. Brain shape convergence in the adaptive radiation of new world monkeys. Proceedings of the National Academy of Sciences 113, 2158–2163 (2016).

[48] Lang, M. M., López-Aguirre, C., Schroeder, L. & Silcox, M. T. Endocranial shape variation and allometry in euarchontoglires. Scientific Reports 14, 17901 (2024).

[49] Boisseau, R. P., Bradler, S. & Emlen, D. J. Divergence time and environmental similarity predict the strength of morphological convergence in stick and leaf insects. Proceedings of the National Academy of Sciences 122, e2319485121 (2025).

[50] Sol, D. et al. Evolutionary divergence in brain size between migratory and resident birds. PLoS One 5, e9617 (2010).

[51] Melchionna, M. et al. Cortical areas associated to higher cognition drove primate brain evolution. Communications biology 8, 80 (2025).

[52] Garin, C. M. et al. An evolutionary gap in primate default mode network organization. Cell reports 39 (2022).

[53] Sokolowski, K. & Corbin, J. G. Wired for behaviors: from development to function of innate limbic system circuitry. Frontiers in molecular neuroscience 5, 55 (2012).

[54] Gutierrez-Barragan, D., Ramirez, J. S., Panzeri, S., Xu, T. & Gozzi, A. Evolutionarily conserved fmri network dynamics in the mouse, macaque, and human brain. Nature Communications 15, 8518 (2024).

[55] Buckner, R. L., Andrews-Hanna, J. R. & Schacter, D. L. The brain’s default network: anatomy, function, and relevance to disease. Annals of the new York Academy of Sciences 1124, 1–38 (2008).

[56] Margulies, D. S. & Smallwood, J. Converging evidence for the role of transmodal cortex in cognition. Proceedings of the National Academy of Sciences 114, 12641–12643 (2017).

[57] Liu, Y. et al. Organization of corticocortical and thalamocortical top-down inputs in the primary visual cortex. Nature Communications 15, 4495 (2024).

[58] Yang, W., Tipparaju, S. L., Chen, G. & Li, N. Thalamus-driven functional populations in frontal cortex support decision-making. Nature neuroscience 25, 1339–1352 (2022).

[59] Mao, T. et al. Long-range neuronal circuits underlying the interaction between sensory and motor cortex. neuron 72, 111–123 (2011).

[60] Barbas, H. General cortical and special prefrontal connections: principles from structure to function. Annual review of neuroscience 38, 269–289 (2015).

[61] Bullmore, E. & Sporns, O. The economy of brain network organization. Nature reviews neuroscience 13, 336–349 (2012).

[62] Humphries, M. D. Dynamical networks: Finding, measuring, and tracking neural population activity using network science. Network Neuroscience 1, 324–338 (2017).

[63] Hynes, R. O. & Zhao, Q. The evolution of cell adhesion. The Journal of cell biology 150, F89–F96 (2000).

[64] Luppi, A. I. et al. Oxygen and the spark of human brain evolution: complex interactions of metabolism and cortical expansion across development and evolution. The Neuroscientist 30, 173–198 (2024).

[65] Miller, A. H., Haroon, E. & Felger, J. C. The immunology of behavior—exploring the role of the immune system in brain health and illness. Neuropsychopharmacology 42, 1–4 (2017).

[66] Castellani, G., Croese, T., Peralta Ramos, J. M. & Schwartz, M. Transforming the understanding of brain immunity. Science 380, eabo7649 (2023).

[67] Shen, Y. et al. Multimodal nature of the single-cell primate brain atlas: Morphology, transcriptome, electrophysiology, and connectivity. Neuroscience Bulletin 40, 517–532 (2024).

[68] Jorstad, N. L. et al. Comparative transcriptomics reveals human-specific cortical features. Science 382, eade9516 (2023).

[69] Adesnik, H. & Naka, A. Cracking the function of layers in the sensory cortex. Neuron 100, 1028–1043 (2018).

[70] Hodge, R. D. et al. Conserved cell types with divergent features in human versus mouse cortex. Nature 573, 61–68 (2019).

[71] Schwartz, E. et al. Evolution of cortical geometry and its link to function, behaviour and ecology. Nature communications 14, 2252 (2023).

[72] Arnatkeviciūtė, A., Fulcher, B. D. & Fornito, A. A practical guide to linking brain-wide gene expression and neuroimaging data. Neuroimage 189, 353–367 (2019).

[73] Jorstad, N. L. et al. Transcriptomic cytoarchitecture reveals principles of human neocortex organization. Science 382, eadf6812 (2023).

[74] Wagstyl, K. et al. Bigbrain 3d atlas of cortical layers: Cortical and laminar thickness gradients diverge in sensory and motor cortices. PLoS biology 18, e3000678 (2020).

[75] Barber, C. B., Dobkin, D. P. & Huhdanpaa, H. The quickhull algorithm for convex hulls. ACM Transactions on Mathematical Software (TOMS) 22, 469–483 (1996).

[76] Yarkoni et al. Large-scale automated synthesis of human functional neuroimaging data. Nature Methods (2011).

[77] Waehnert, M. et al. Anatomically motivated modeling of cortical laminae. Neuroimage 93, 210–220 (2014).

[78] Alexander et al. Local-global parcellation of the human cerebral cortex from intrinsic functional connectivity mri. Cerebral Cortex (2017).

[79] Wu, T. et al. clusterprofiler 4.0: A universal enrichment tool for interpreting omics data. The innovation 2 (2021).

