## Supplemental Information for "Sulcal landmarks reveal lineage-specific trajectories and multi-scale specialization of the mammalian cortex"

### Supplementary Figures

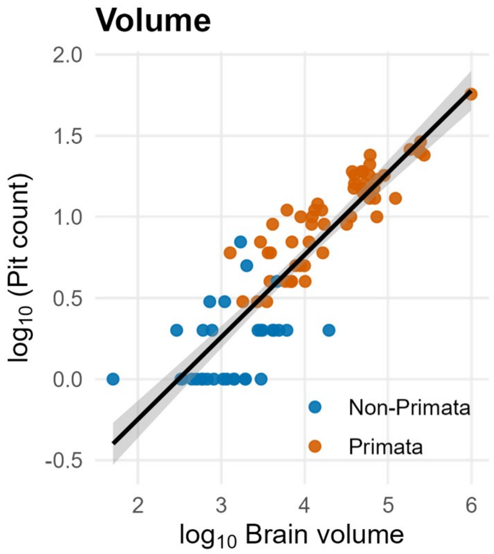


Supplementary Figure 1: Log–log scaling of pit count with brain volume. Scatterplot of $\log_{10}$(pit count) versus $\log_{10} V$ (brain volume), with an OLS fit and 95% confidence band; points are colored by primate vs. non-primate. The slope is $\hat{\beta}_V = 0.507 \pm 0.027$ ($t = 18.46$, $p < 2 \times 10^{-16}$), explaining $R^2 = 0.795$ of the variance (residual s.e. = 0.217). A tenfold increase in $V$ corresponds to $\sim 10^{0.507} \approx 3.2\times$ more pits, consistent with sublinear scaling.

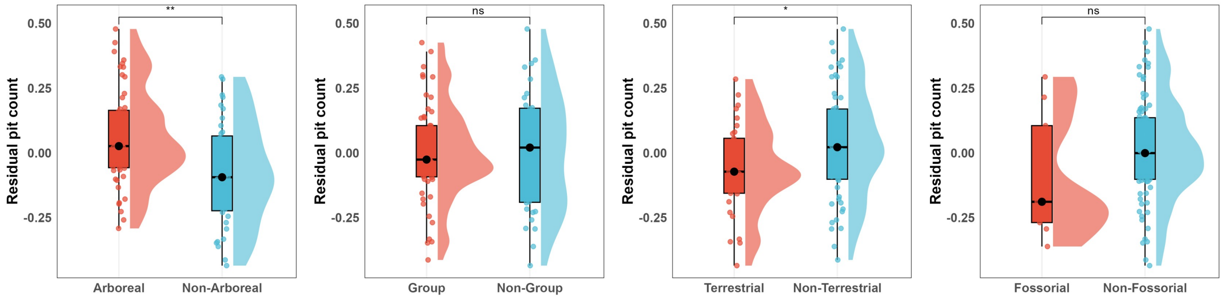


Supplementary Figure 2: Size-adjusted residual pit counts across ecological categories. Vertical raincloud plots compare Arboreal vs. Non-Arboreal, Group vs. Non-Group, Terrestrial vs. Non-Terrestrial, and Fossorial vs. Non-Fossorial. Residual pit counts were obtained from a multivariable log–linear model jointly controlling for cortical surface area and brain volume. Significance (two-sided Wilcoxon, FDR-corrected) is annotated above each contrast: $*$ indicates $p < 0.05$ and $**$ indicates $p < 0.01$.

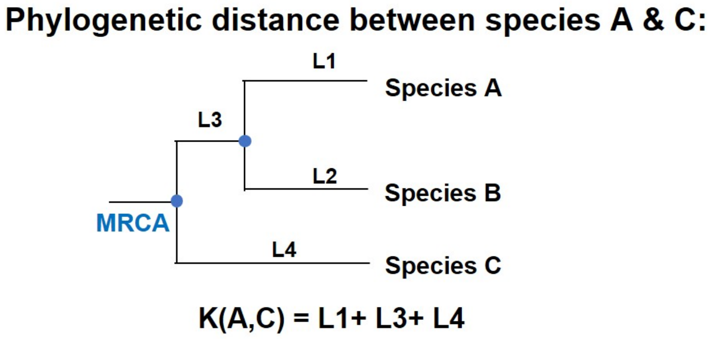


Supplementary Figure 3: Method for computing the phylogenetic distance between species A and C. On a time-calibrated phylogeny, the distance is defined as the total branch length along the shortest path linking A and C. Operationally, identify their most recent common ancestor (MRCA) and sum the branch lengths from each tip to the MRCA: $K(A, C) = \mathrm{L1} + \mathrm{L2} + \mathrm{L3}$. Branch lengths are measured in millions of years (Myr).

**(A) Pit Evolution in Non-Primate**

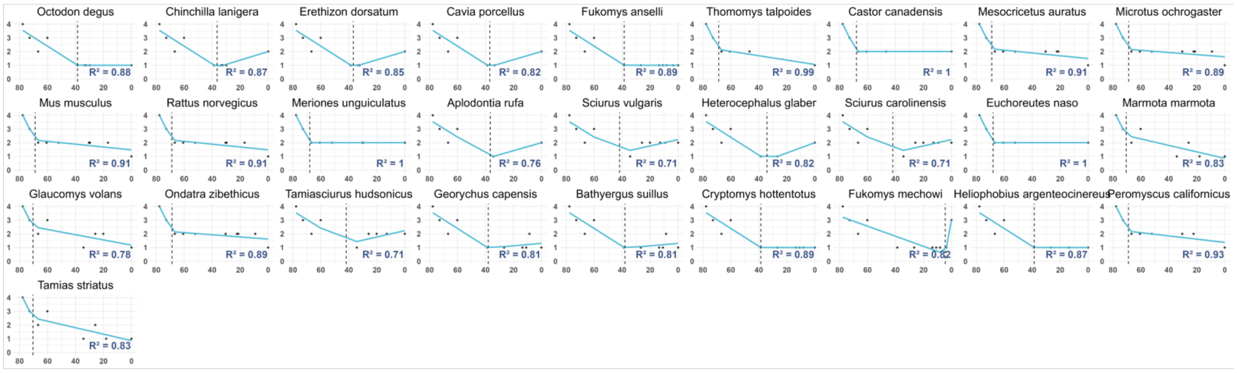


**(B) Pit Evolution in Primate**

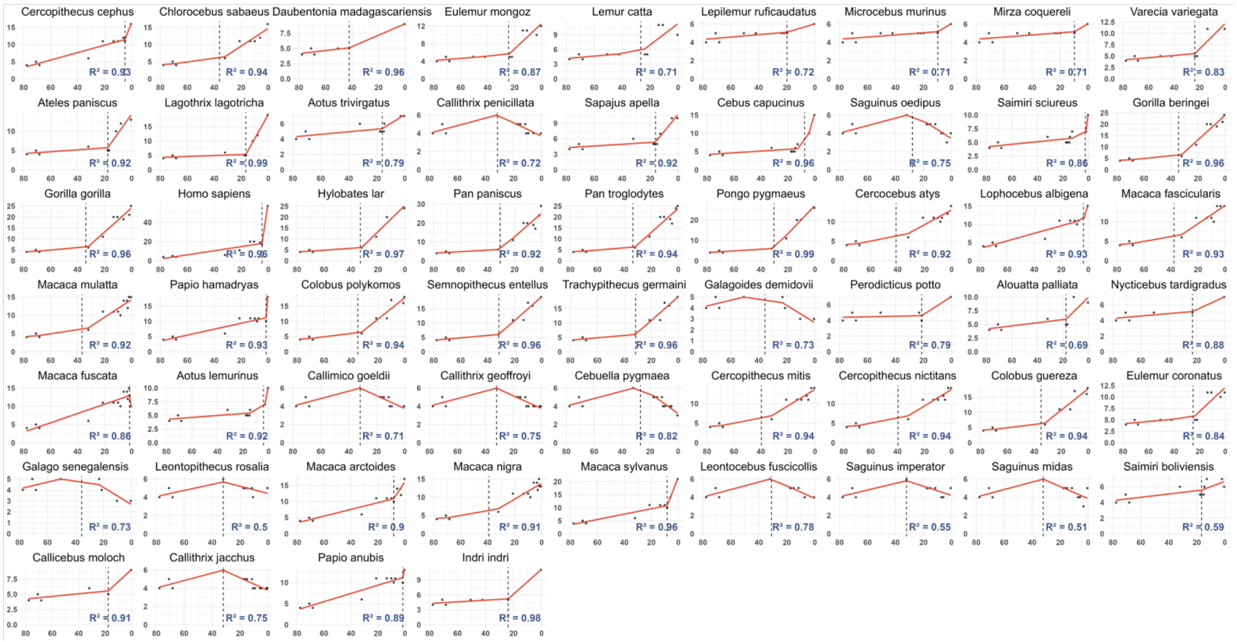


Supplementary Figure 4: Evolutionary trajectories of sulcal pit counts across 90 extant species. For each species, a piecewise linear regression was fitted to the ancestor-to-present time series; $R^2$ denotes goodness of fit and the breakpoint marks the evolutionary turning point. Most primate trajectories remain flat through deep time and then rise sharply over the last 20 Mya, consistent with late-phase acceleration. In contrast, most non-primate trajectories show an early decline ( 80–30 Mya) followed by a prolonged plateau, yielding a more stable, flattened pattern. To ensure accurate and meaningful piecewise fits, only lineages with $> 3$ dated nodes between 80 Mya and the present were included; four species did not meet this criterion and are not shown. (A) 28 non-primate species; (B) 58 primate species.

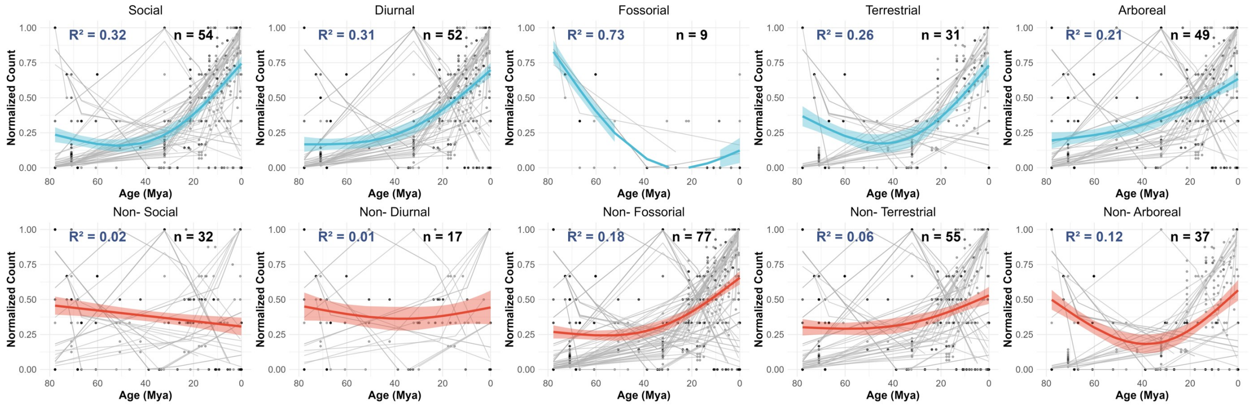


Supplementary Figure 5: For each species, pit-count trajectories were normalized to $[0, 1]$ and fitted with a GAM ($k = 3$); panels display the group-level fit (colored curve) with 95% confidence band overlaid on species-wise trajectories (gray), with the number of species ($n$) and GAM $R^2$ reported in each panel. Contrasts are shown for Social vs. Non-Social, Diurnal vs. Non-Diurnal, Fossorial vs. Non-Fossorial, Terrestrial vs. Non-Terrestrial, and Arboreal vs. Non-Arboreal groups. These summaries reveal distinct evolutionary tempos across categories, indicating that *pit-based* evolutionary trajectories sensitively reflect ecology- and lifestyle-related differences across species.

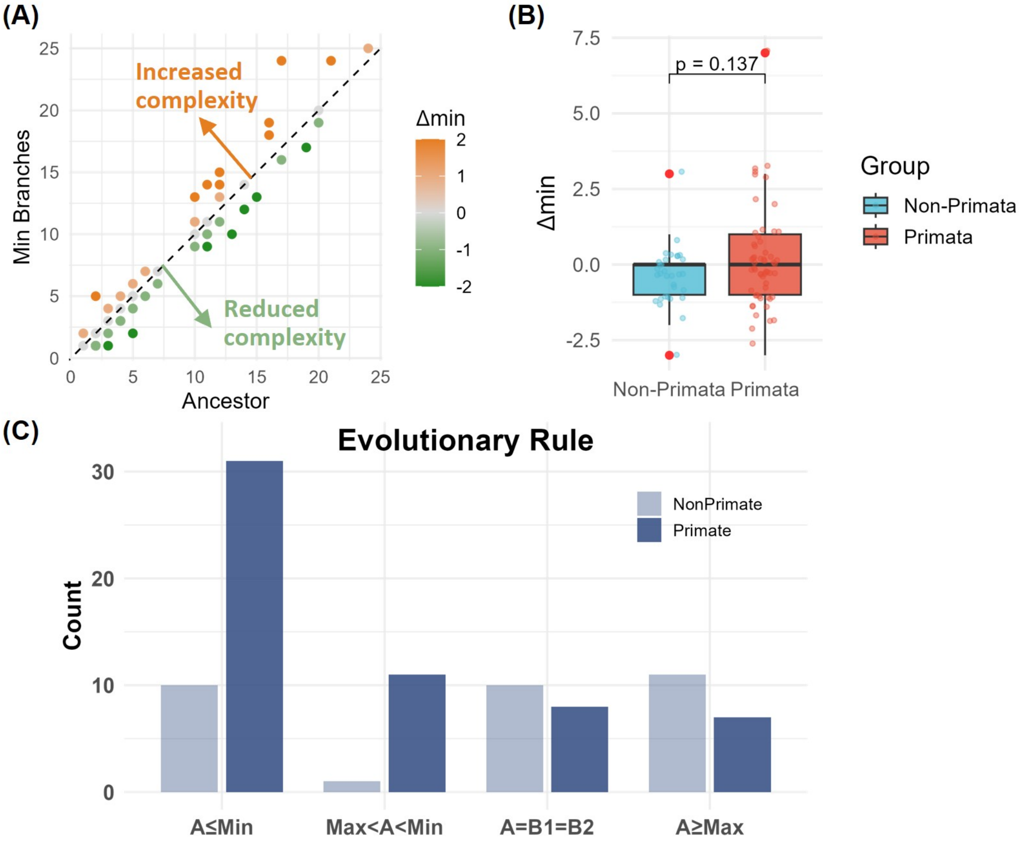


Supplementary Figure 6: Post-split change in pit-defined folding using a conservative 'minimum descendant' criterion. For each internal split we compute $\Delta_{\min} = \min(B_1, B_2) - A$, where $A$ is the ancestor's whole-brain pit count and $B_1, B_2$ are the two descendants. (A) Scatter of descendant minimum vs. ancestor with the $y = x$ line; points above indicate that both descendants have higher pit counts than the ancestor (increased folding complexity), whereas points below indicate reductions. (B) Boxplots of $\Delta_{\min}$ by group. although the difference is not significant (two-sided Wilcoxon $p = 0.137$), the median in primates is higher than in non-primates. (C) counts of split types classified by the ancestor–descendant relation: (i) $A \leq \min(B_1, B_2)$, (ii) $\min(B_1, B_2) < A < \max(B_1, B_2)$, (iii) $A = B_1 = B_2$, (iv) $A \geq \max(B_1, B_2)$; primates are enriched in (i), whereas non-primates show more mixed or non-increase cases.

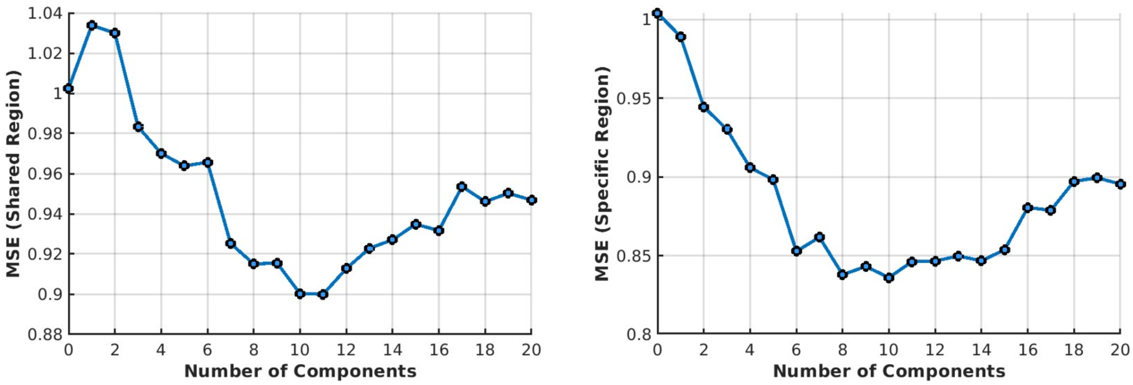


Supplementary Figure 7: Cross-validated selection of the number of PLSR components. For *Homo sapiens*–specific regions and species-shared regions, we performed 10-fold cross-validation and plotted the held-out mean squared error (MSE) as a function of the number of components. In both analyses the MSE reached its minimum at $h = 10$, which was therefore used in the final PLSR reported in the Methods. The curves show the typical U-shape: error decreases with added components and rises again beyond the optimum.

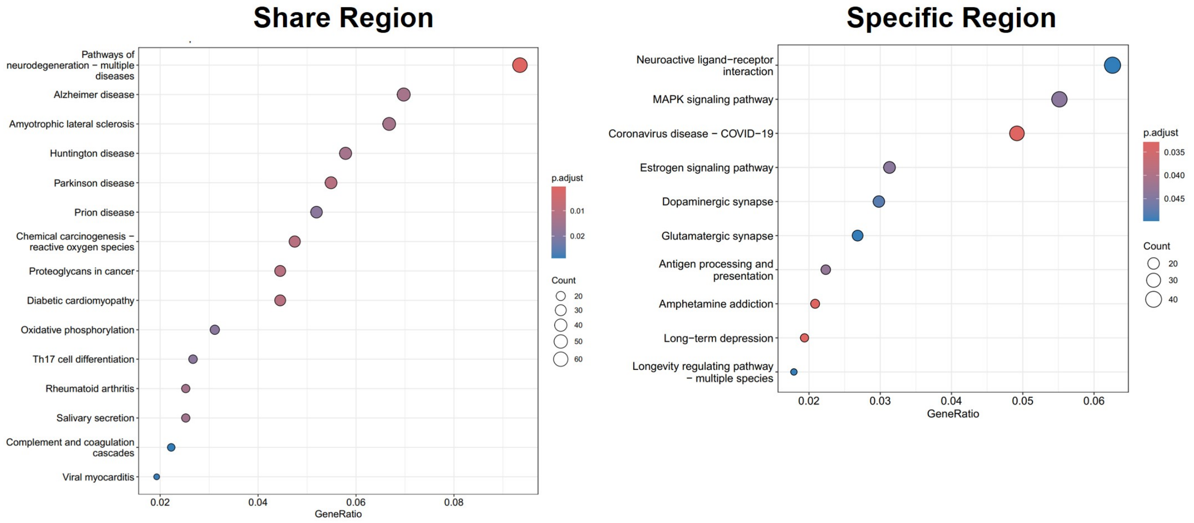


Supplementary Figure 8: KEGG pathway enrichment for *Homo sapiens*-specific and species-shared cortical regions. Bubble plots show the top enriched KEGG pathways for genes associated with species-shared regions (left) and *Homo sapiens*-specific regions (right). The x-axis represents the gene ratio (proportion of genes in each pathway), bubble size indicates the number of genes, and bubble color denotes adjusted p values (FDR-corrected). Shared regions are mainly enriched in pathways related to neurodegenerative and metabolic diseases, whereas *Homo*-specific regions are enriched in signaling and synaptic transmission pathways.

### Supplementary Tables

Supplementary Table 1: Results of multiple regression on distance matrices (MRM). Standardized coefficients (std) and raw-scale coefficients (raw) are reported, together with permutation-based $p$ values (10,000 permutations). Phylogenetic distance remained significant in both models, whereas cortical surface area and brain volume differences were not significant.

| **Term** | **Beta (std)** | $p_{\mathbf{perm}}$ **(std)** | **Beta (raw)** | $p_{\mathbf{perm}}$ **(raw)** |
|---|---|---|---|---|
| Intercept | 0.519 | 1.000 | 0.514 | 1.000 |
| Phylogenetic distance | 0.136 | $< 0.001$ | 0.136 | $< 0.001$ |
| Surface area difference | 0.048 | 0.418 | $1.53 \times 10^{-6}$ | 0.418 |
| Brain volume difference | $-0.046$ | 0.446 | $-3.05 \times 10^{-7}$ | 0.446 |
| **Model** $R^2$ | 0.124 | $< 0.001$ | 0.124 | $< 0.001$ |
